# Mechanochemically-reprogrammed stem cell exosomes reconcile the biogenesis internalization trade-off for pulmonary fibrosis therapy

**DOI:** 10.64898/2026.03.20.713315

**Authors:** Chenxi Pan, Chuanfeng An, Zhen He, Kaiwen Chen, Yi He, Yonggang Zhang, Tengfei Tian, Xiangying Wang, Huanan Wang

## Abstract

Matrix stiffness serves as a pivotal biophysical cue of extracellular vesicle (EV) function and cellular uptake, yet the mechanical cues that enhance EV function may not simultaneously favour cellular uptake. Through comparative analyses across pulmonary cell populations, we identified macrophages as the predominant EV recipients in our experimental system and as a functionally relevant effector population in pulmonary fibrosis. Herein, we developed a mechanochemical reprogramming strategy to coordinate the immunomodulatory function of mesenchymal stem cell (MSC)-derived EVs with their uptake by macrophages. We found that soft-matrix conditioning of MSC-derived EVs enhanced immunomodulatory activity and promoted an anti-inflammatory macrophage phenotype. By contrast, increasing matrix stiffness upregulated ATP-binding cassette transporter A1 (ABCA1), increased EV membrane cholesterol, and enhanced macrophage uptake. Guided by these complementary mechanical responses, we combined soft-matrix conditioning with ABCA1 modulation to generate mechanochemically programmed MSC-EVs. This strategy preserved the immunomodulatory advantages associated with the soft matrix while enhancing macrophage uptake and increasing EV production. In a murine model of pulmonary fibrosis, mechanochemically reprogrammed EVs produced higher and more persistent EV-associated signals in lung tissue and were preferentially associated with pulmonary macrophages. Treatment promoted anti-inflammatory and reparative macrophage responses and was accompanied by reduced pulmonary inflammation and fibrotic remodelling. Together, these findings establish a mechanochemically reprogrammed strategy that coordinates MSC-EV immunomodulatory function with macrophage uptake while increasing EV production, providing a framework for engineering EV-based therapies for pulmonary fibrosis.

## 1. Introduction

Extracellular vesicles (EVs) have emerged as a promising cell-free platform for harnessing the therapeutic activity of mesenchymal stem cells (MSCs) [1–3]. EVs can transport bioactive proteins, lipids and nucleic acids while retaining favourable biocompatibility and stability in biological fluids [4]. Among their proposed therapeutic activities, the immunomodulatory effects of MSC-derived EVs are particularly relevant to inflammatory and fibrotic diseases, in which dysregulated immune responses contribute substantially to disease progression [5]. Despite this potential, several barriers continue to limit the translation of MSC-derived EVs. EV production under conventional culture conditions is relatively low, limiting scalable manufacturing and increasing production costs [6]. In addition, therapeutic efficacy depends not only on EV cargo but also on efficient delivery to relevant recipient cells. After administration, the mononuclear phagocyte system can rapidly clear EVs, limiting bioavailability and causing variable tissue exposure [7]. Rather than simply increasing EV production, future engineering strategies should prioritize improving EV therapeutic quality and their effective utilization by disease-relevant recipient cells.

Matrix stiffness offers a bioinspired and experimentally reproducible way to engineer EVs, as it recapitulates a fundamental biophysical cue of the native cellular microenvironment [8]. Previous studies have shown that stiffness-tunable hydrogels can mechanically condition stem cells and alter the properties of the EVs they release, with softer matrices often favouring EVs with enhanced anti-inflammatory activity [9]. These stiffness-dependent effects are closely associated with cytoskeletal remodelling and accompanying changes in cell morphology. Cells cultured on soft matrices typically adopt a more rounded morphology, with reduced spreading and increased local membrane curvature, whereas cells on stiffer matrices become more spread and elongated [10]. Such changes in cell shape, membrane tension and cortical actin organization can influence membrane remodelling and vesicular trafficking. In particular, increased membrane curvature may lower the energetic cost of local membrane deformation and budding, thereby facilitating the formation and release of specific EV populations [11]. Together, these mechanically regulated processes provide a mechanistic basis for engineering EV production and function by controlling the cellular microenvironment.

Recent studies have shown that EVs generated under soft-matrix conditions can exhibit higher production while showing lower delivery efficiency than those derived from stiffer matrices [10]. However, the molecular mechanisms underlying this stiffness-dependent difference in cellular uptake remain poorly understood. As a result, strategies based solely on physical modulation of matrix stiffness may improve one aspect of EV performance while compromising another, limiting their broader engineering potential [10, 12]. Identifying the regulatory nodes that uncouple these stiffness-responsive responses could therefore provide a rational basis for engineering EVs with improved functional quality, delivery efficiency and manufacturing performance.

Herein, we developed a mechanochemical reprogrammed strategy that combines soft-matrix conditioning with chemical modulation of ATP-binding cassette transporter A1 (ABCA1) to enhance MSC-derived EV production and macrophage uptake while preserving their immunomodulatory activity (Figure 1). Comparative analyses across multiple lung-associated cell populations identified macrophages as the predominant EV recipients in our experimental system and as a major functional effector population in pulmonary fibrosis. By integrating transcriptomic profiling with biomechanical analyses, we further identified ABCA1 as a mechanosensitive regulator linking matrix stiffness to changes in EV membrane cholesterol and established membrane cholesterol as an important determinant of EV uptake. We therefore focused on how mechanically regulated membrane remodelling shapes EV-macrophage interactions. Guided by these findings, chemical activation of ABCA1 in MSCs cultured on soft matrices enriched EV membrane cholesterol and partially recapitulated the uptake advantage associated with stiffer matrices, while retaining the immunomodulatory properties conferred by the soft matrices and further increasing EV production. Together, these findings establish a framework for EV engineering that coordinates functional quality, cellular delivery and production, providing an engineering basis for developing more effective and scalable EV-based therapies.

**Figure 1.**
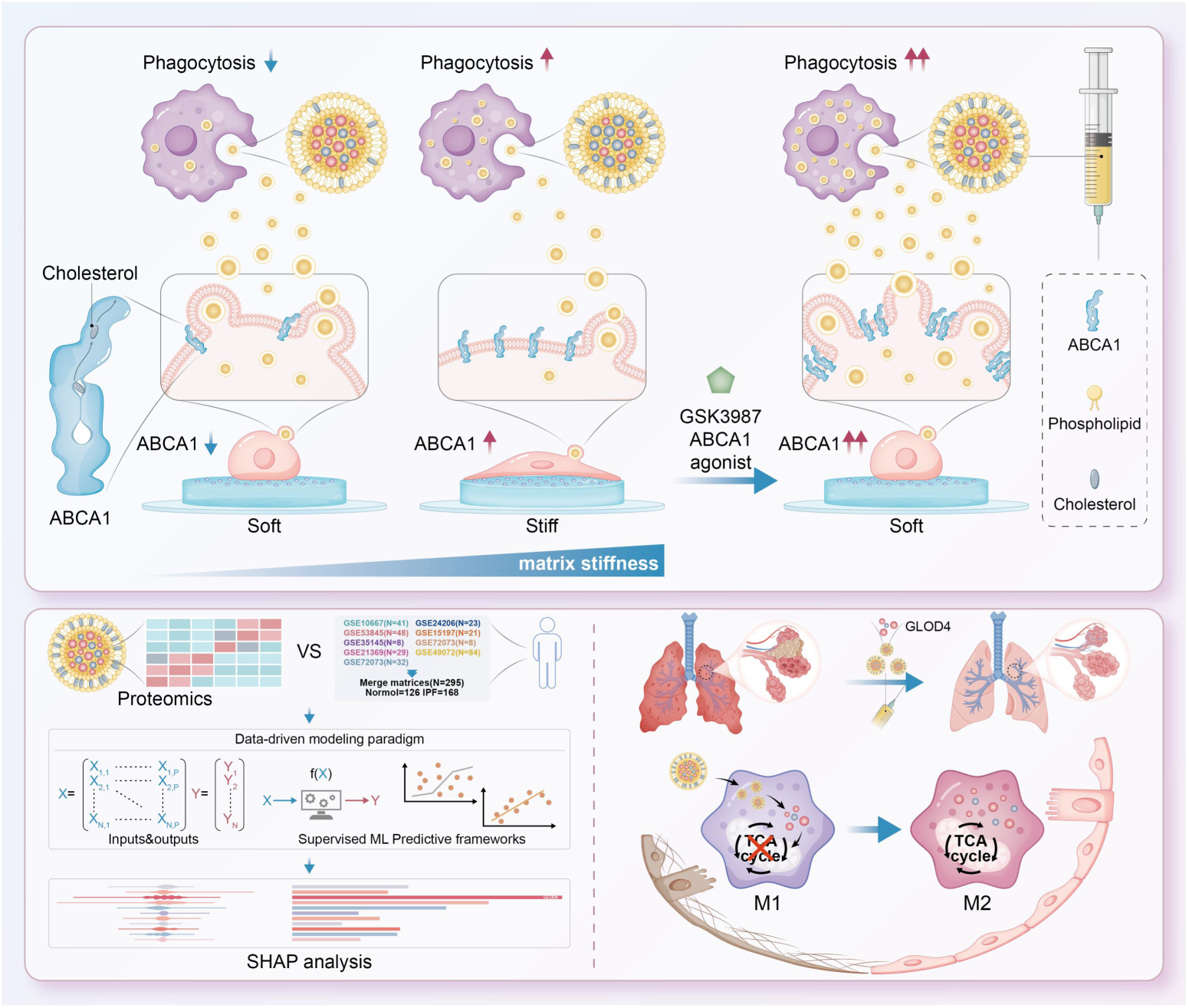
Schematic illustration of a mechanochemically reprogramming strategy for generating therapeutic MSC-derived EVs, with macrophages highlighted as the central effector cells mediating their anti-inflammatory activity.

## 2. Results

### 2.1 Identification of ABCA1 as a mechanosensitive regulator linking matrix stiffness to lipid transport

Previous studies of MSC mechanosensitivity have largely focused on changes in cell morphology and cytoskeletal organization. To investigate the transcriptional programmes associated with these mechanical responses, we analysed the GSE22641 dataset, which used a GelMA hydrogel system with stiffness values comparable to those in our study (∼3 kPa and ∼30 kPa) [13]. Using a similar material system allowed us to compare stiffness-dependent transcriptional responses while minimizing differences arising from material composition [14]. Differential expression analysis identified four genes that were consistently upregulated under the stiffer matrix condition: Synaptopodin 2 (SYNPO2), ATP-binding cassette transporter A1 (ABCA1), platelet-derived growth factor D (PDGFD), and Rho GTPase activating protein 28 (ARHGAP28) (Figure 2A). Volcano plot analysis further showed that these candidates were significantly associated with matrix stiffness (Figure 2B).

**Figure 2.**
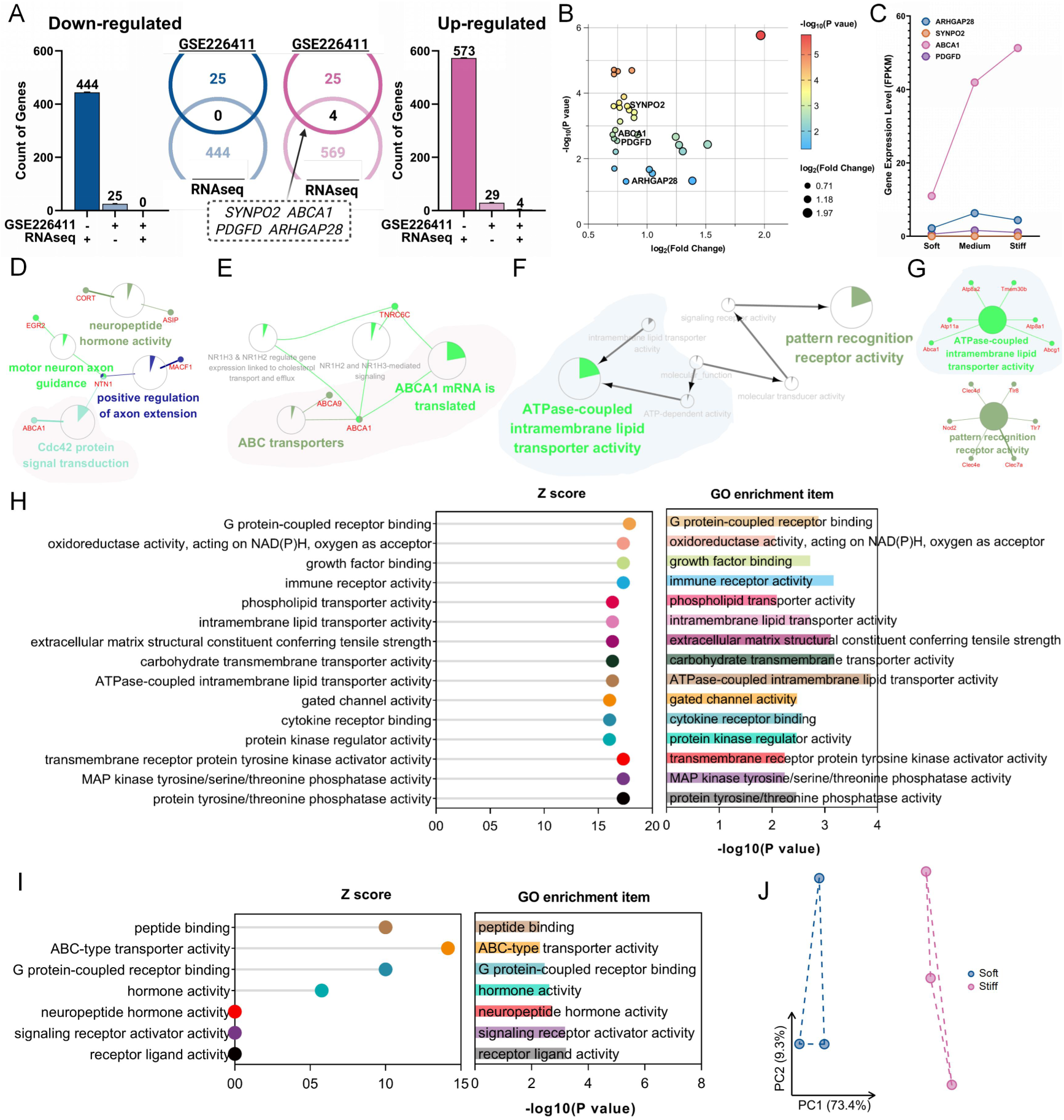
Identification of ABCA1 as a mechanosensitive regulator through integrative transcriptomic analysis. (A) Venn diagram showing the overlap between differentially expressed genes identified in our RNA-seq dataset and the public GSE22641 dataset, yielding four candidate genes shared by both datasets: SYNPO2, ABCA1, PDGFD and ARHGAP28. (B) Volcano plot of differentially expressed genes in the GSE22641 dataset, with the four overlapping candidates highlighted. (C) Expression profiles of SYNPO2, ABCA1, PDGFD and ARHGAP28 across soft, intermediate and stiff matrix conditions, showing a monotonic increase in ABCA1 expression with increasing matrix stiffness. (D, E) KEGG and Gene Ontology enrichment analyses of differentially expressed genes in the GSE22641 discovery dataset. (F, G) KEGG and Gene Ontology enrichment analyses of differentially expressed genes in the independent validation dataset. (H, I) Z-score normalized comparison of Gene Ontology enrichment patterns across the two datasets, highlighting stiffness-responsive changes in ABCA1-related lipid-transport pathways. (J) Principal component analysis based on genes associated with the ABCA1-related lipid-transport programme, showing separation of samples cultured under soft and stiff matrix conditions (GSE22641 dataset).

To further define the stiffness-dependent expression patterns of these candidates, we examined their expression across a matrix-stiffness gradient of 3.4, 14.2 and 31.4 kPa. We reasoned that candidates showing a consistent response across this gradient would be more likely to participate in stiffness-dependent regulation [14]. Among the four genes, ABCA1 showed the clearest monotonic response. Although SYNPO2 also varied with matrix stiffness, its overall expression remained very low (FPKM < 0.1). By contrast, ABCA1 was expressed at relatively high levels, increasing approximately 5-fold from the softest to the stiffest matrices. PDGFD and ARHGAP28 showed non-monotonic responses, with higher expression at intermediate stiffness values (Figure 2C). Its pronounced stiffness responsiveness and consistent monotonic expression across the stiffness gradient therefore led us to prioritize ABCA1 for subsequent mechanistic investigation.

To further evaluate the functional significance of the candidate genes, we analysed the GSE22641 dataset, which contains transcriptomic profiles of hMSCs cultured under different matrix-stiffness conditions (Figures 2D, E). KEGG and Gene Ontology analyses identified stiffness-responsive changes in lipid-transport related pathways, and similar enrichment patterns were observed in our independent validation dataset (Figures 2F, G). To quantitatively compare pathway enrichment between the two datasets, we normalized pathway enrichment scores using Z-scores [15]. This analysis consistently highlighted ATP-dependent transmembrane lipid transport, including phospholipid translocation and ATPase-coupled lipid efflux, in which ABCA1 has an established role (Figures 2H, I). We next asked whether expression of genes associated with the ABCA1-related lipid-transport programme could distinguish cells cultured on matrices of tunable stiffness. Principal component analysis of these genes separated samples cultured on soft and stiff matrices into distinct clusters (Figure 2J) [16]. Together, these analyses support a close association between matrix stiffness, ABCA1-related lipid transport and membrane-lipid regulation, providing mechanistic rationale for focusing on ABCA1 in subsequent mechanistic studies.

### 2.2 Validation of the mechanosensitivity of ABCA1 using PAA hydrogels with independently tunable stiffness

To validate ABCA1 mechanosensitivity independently, we established a stiffness-tunable polyacrylamide (PAA) hydrogel platform [17]. Compared with natural hydrogels such as GelMA, the synthetic PAA system allows matrix stiffness to vary while keeping other physicochemical cues comparable, enabling stiffness to be examined as the primary mechanical variable. By adjusting the cross-linker composition, we generated hydrogels with Young’s moduli ranging from 2.62 ± 0.14 to 42.85 ± 2.05 kPa (Figure 3A-E and Figure S1, Supporting Information). To promote MSC adhesion, type I collagen was covalently immobilized onto the hydrogel surfaces. BCA assays and contact-angle measurements showed comparable collagen surface density (∼5 μg cm⁻²) and hydrophilicity/hydrophobicity across the stiffness-tunable platform (Figure S2A-C, Supporting Information). Together, these results established an independent, stiffness-tunable platform for evaluating ABCA1 mechanosensitivity while minimizing variation in material composition and surface properties.

**Figure 3.**
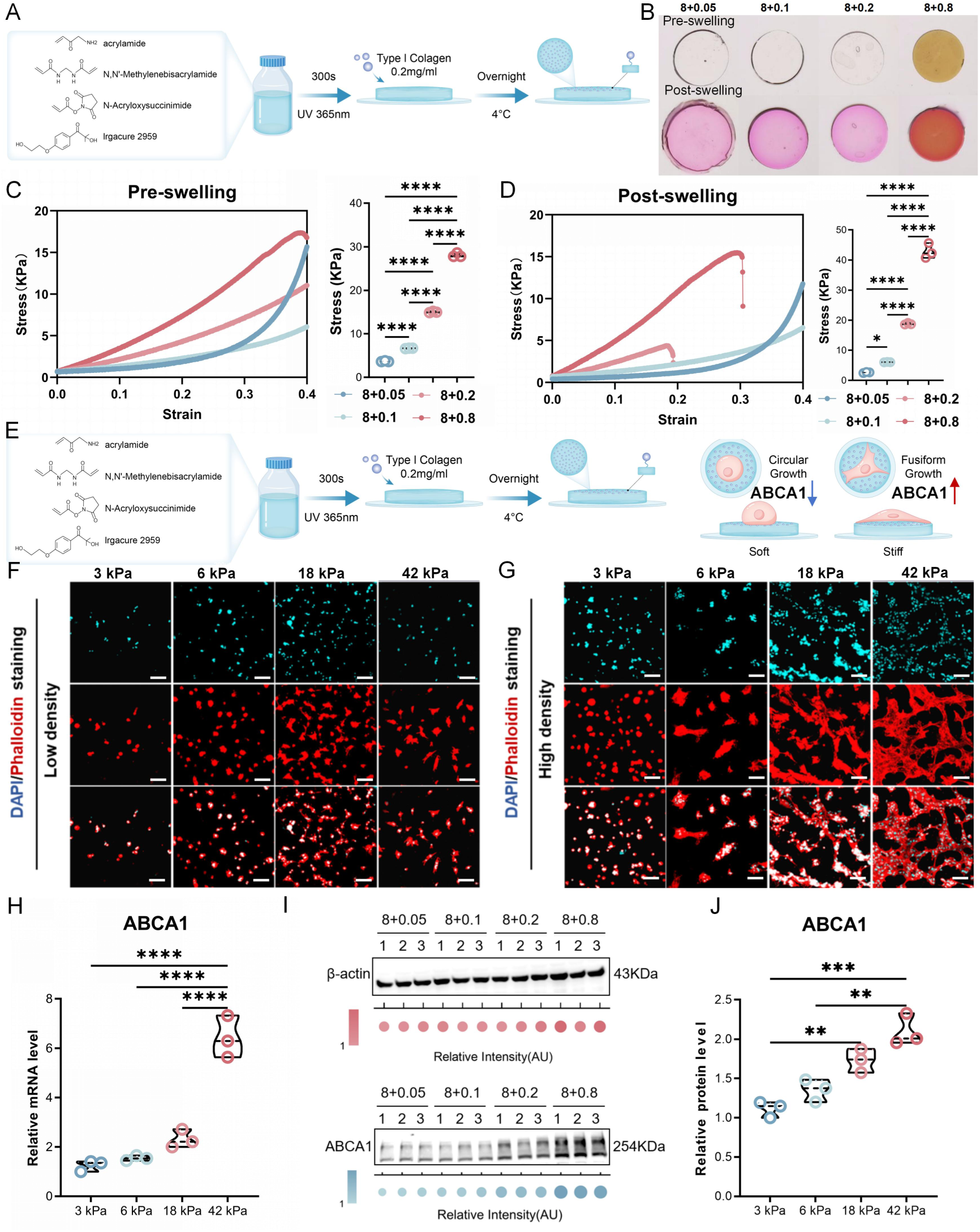
Establishment of a stiffness-tunable PAA hydrogel platform and validation of the mechanosensitivity of ABCA1. (A) Schematic of the composition and fabrication of the stiffness-tunable polyacrylamide (PAA) hydrogel platform. (B) Representative macroscopic images of the four hydrogel formulations before and after equilibrium swelling. (C) Young’s modulus of the hydrogel formulations measured before swelling (n = 3). (D) Young’s modulus of the corresponding hydrogels measured after equilibrium swelling under the indicated conditions (n = 3). (E) Schematic workflow for hydrogel preparation and surface functionalization to promote MSC adhesion. (F) Representative confocal fluorescence images showing MSC morphology and cytoskeletal organization on stiffness-tunable PAA hydrogel platform. (H) RT-qPCR analysis of ABCA1 mRNA expression in MSCs cultured on stiffness-tunable PAA hydrogel platform (n = 3). (I) Representative Western blot analysis of ABCA1 protein expression in MSCs cultured on stiffness-tunable PAA hydrogel platform. (J) Densitometric quantification of ABCA1 protein expression shown in (I) (n = 3).

To determine whether the PAA platform could reliably capture mechanosensitive MSC behaviors, we next evaluated cell viability, proliferation, morphology and YAP localization across the stiffness-tunable platform. Cell viability remained high under all conditions, whereas proliferation and morphology varied with matrix stiffness. MSCs showed limited spreading on softer matrices and progressively greater spreading and elongation on stiffer matrices (Figure 3F and Figure S3, Supporting Information). We further assessed YAP localization as a readout of mechanotransduction. YAP was predominantly cytoplasmic on soft matrices but increasingly localized to the nucleus as matrix stiffness increased (Figure S2D, Supporting Information) [17]. These results support using the PAA platform for subsequent analysis of the mechanosensitivity of ABCA1

To determine whether the mechanosensitivity of ABCA1 response identified by transcriptomic screening was reproduced in the independent PAA system, we quantified ABCA1 expression across the stiffness-tunable platform by RT-qPCR and Western blotting. Both analyses showed that ABCA1 expression increased with matrix stiffness (Figure 3H-J), consistent with the transcriptional pattern observed in the GelMA-based dataset. This cross-platform agreement supports ABCA1 as a reproducible mechanosensitive regulator across distinct material systems, while minimizing the likelihood that the observed response was specific to a particular hydrogel chemistry.

The human ABC transporter superfamily comprises 49 members, classified into seven subfamilies (A-G) [18]. While these transporters handle diverse substrates, subfamilies A and G are pivotal for translocating lipids across the plasma membrane [19]. Specifically, ABCA1 orchestrates the crosstalk between lipid metabolism and EV biogenesis by driving cholesterol efflux, a process essential for EV release [20]. In this context, EVs function as key delivery vehicles that reportedly account for approximately 30% of intercellular cholesterol transport [21, 22]. Building on evidence that mechanical cues from cell-matrix adhesion can induce lipid metabolic shifts in MSCs to fuel neutral lipid and cholesterol synthesis [23], we postulated that matrix stiffness dictates the cholesterol content of MSC-derived EVs through an ABCA1-mediated mechanism.

### 2.3 Differential regulation of EV cargo potency and macrophage-targeted delivery efficiency by matrix stiffness

To examine whether matrix stiffness regulates the membrane cholesterol content of MSC-derived EVs through ABCA1, we next used the stiffness-tunable PAA platform described above. MSCs were cultured on PAA hydrogels spanning the defined stiffness range, then EVs were isolated from the conditioned medium for downstream characterization and functional analyses (Figure 4A). This experimental framework enabled us to directly assess how stiffness-dependent changes in ABCA1 expression levels are reflected in EV physicochemical properties, membrane composition, and interactions with multiple recipient cell types.

**Figure 4.**
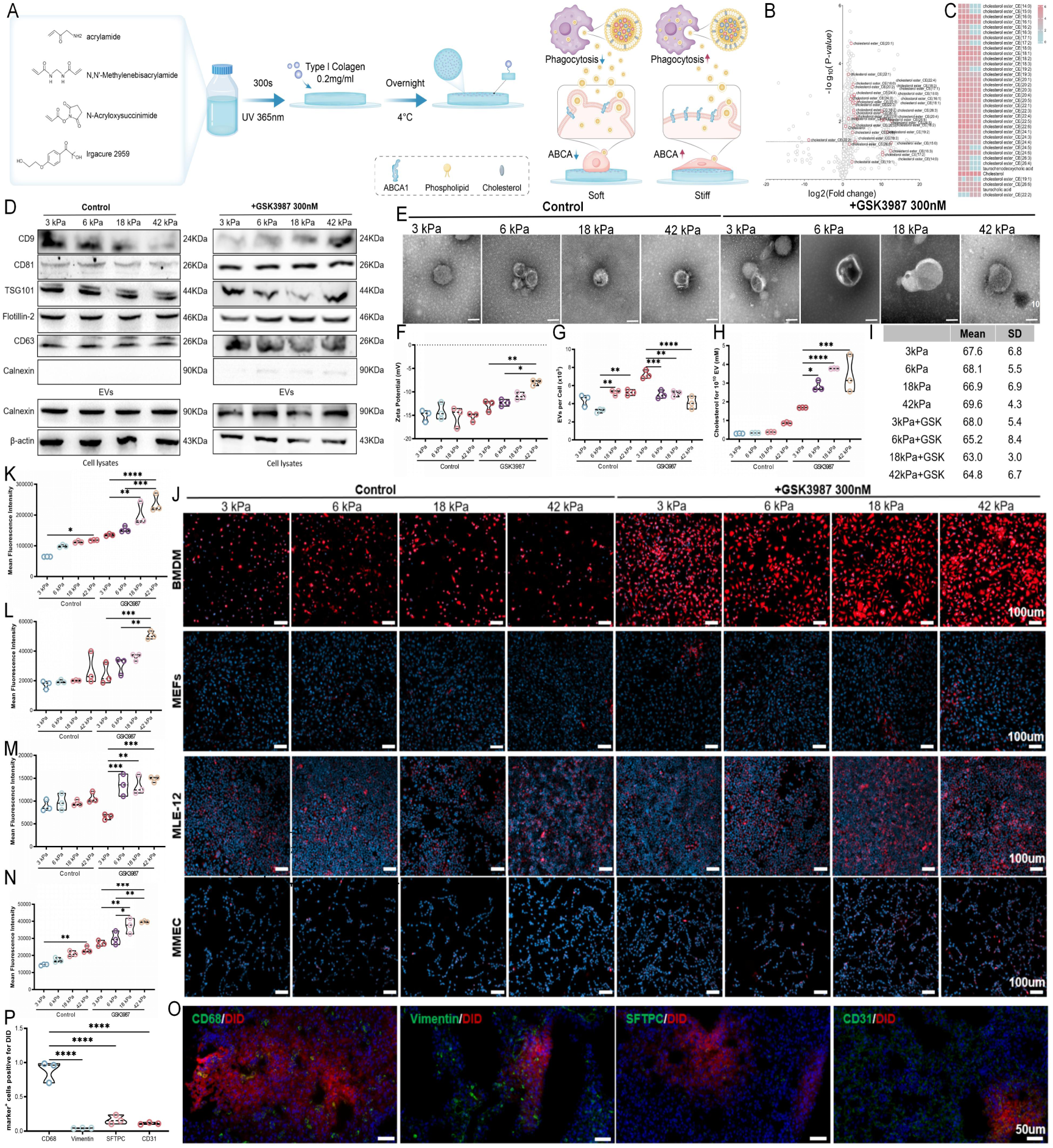
Macrophages are the predominant recipients of MSC-derived EVs with uptake regulated by matrix stiffness. (A) Schematic of the experimental workflow. MSCs were cultured on a stiffness-tunable PAA hydrogel platform, and the resulting EVs were collected for physicochemical characterization, membrane cholesterol analysis, and recipient cell uptake assays. (B) Volcano plot of differential metabolomic profiles between stiffness-tunable conditions, highlighting cholesterol-related metabolites associated with matrix stiffness. (C) Heatmap of cholesterol-related metabolic profiles under stiffness-tunable conditions. (D) Western blot analysis of EV-associated markers in MSC-derived EVs generated under stiffness-tunable conditions. (E) Representative transmission electron microscopy (TEM) images of MSC-derived EVs generated under stiffness-tunable conditions. (Scale bar = 100 nm). (F) Zeta potential of MSC-derived EVs generated under stiffness-tunable conditions (n = 3). (G) Nanoflow cytometric quantification of EV particle numbers produced by MSCs under the indicated mechanical and chemical conditions (n = 3). (H) Membrane cholesterol content of MSC-derived EVs generated under the indicated mechanical and chemical conditions (n = 3). (I) Particle-size distributions (nm) of MSC-derived EVs generated under stiffness-tunable conditions. (J) Representative confocal fluorescence images showing uptake of equal numbers of DiD-labelled EV particles by bone marrow-derived macrophages (BMDMs), mouse embryonic fibroblasts (MEFs), MLE-12 pulmonary epithelial cells and mouse microvascular endothelial cells (MMECs). (K-N) Quantification of fluorescence associated with DiD-labelled EV uptake by BMDMs (K) (n = 3), MEFs (L) (n = 3), MLE-12 cells (M) (n = 3) and MMECs (N) (n = 3). (O) Representative fluorescence images showing the cellular distribution of DiD-labelled EV fluorescence in pulmonary fibrosis. Seven days after induction of pulmonary fibrosis, mice received DiD-labelled mechanochemically reprogrammed EVs generated from MSCs cultured on soft matrices with chemical activation of ABCA1 by GSK3987, and lung tissues were collected 48 h later. CD68, Vimentin, SFTPC and CD31 were used to identify macrophages, fibroblasts, pulmonary epithelial cells and microvascular endothelial cells, respectively. (P) Quantification of colocalization between DiD-labelled EV fluorescence and CD68⁺ macrophages, Vimentin⁺ fibroblasts, SFTPC⁺ pulmonary epithelial cells and CD31⁺ microvascular endothelial cells (n = 3).

To systematically characterize the effects of matrix stiffness on the physicochemical properties of MSC-derived EVs, EVs collected from MSCs cultured on matrices of different stiffness were analyzed by Western blotting, transmission electron microscopy (TEM), zeta-potential measurement and nanoflow cytometry. Western blotting confirmed the expression of canonical EV markers, including CD9, CD81, TSG101, Flotillin-2 and CD63, together with the absence of the endoplasmic reticulum marker Calnexin, supporting the purity of the isolated EVs (Figure 4D). MSC-EVs obtained under different stiffness conditions exhibited comparable negative zeta potentials of approximately -12.99 ± 2.69 mV (Figure 4F). TEM and nanoflow cytometry further showed similar size distributions across groups, with mean EV diameters ranging from approximately 63 to 69 nm (Figures 4E and I). Quantitative nanoflow cytometry showed no significant difference in EV particle numbers between the soft- and stiff-matrix conditions when EVs were enriched by ultrafiltration (Figure 4H). The absence of a detectable difference may partly reflect technical variability associated with the ultrafiltration procedure, including membrane adsorption and incomplete EV recovery, which could reduce sensitivity to modest differences in EV yield.

To determine an appropriate GSK3987 concentration for subsequent mechanochemical programming, we performed a dose-response analysis in MSCs and evaluated both ABCA1 expression and cholesterol distribution between the cells and culture medium [24]. Because GSK3987 treatment aimed to enhance ABCA1-dependent cholesterol efflux, we selected the treatment condition based on functional cholesterol mobilization rather than ABCA1 expression alone. Cholesterol levels in the culture supernatant increased with GSK3987 treatment and peaked at 300 nM, accompanied by the lowest cellular cholesterol content (Figure S4, Supporting Information). Although ABCA1 expression changed further at 500 nM, it was not accompanied by a corresponding improvement in cholesterol efflux. We therefore selected 300 nM GSK3987 for subsequent experiments as the condition that produced the most favourable cholesterol-efflux phenotype.

We next investigated whether chemical activation of ABCA1 could further influence EV production. Under the soft-matrix condition, GSK3987 treatment increased EV particle numbers to nearly twice those produced by untreated MSCs cultured on the same matrix (Figure 4H). Chemical activation of ABCA1 therefore enhanced EV production under the soft-matrix condition. Given this increase in EV yield, we next examined whether GSK3987 treatment also altered the functional properties of the resulting EVs.

Before examining the downstream functional effects of MSC-derived EVs, we first sought to identify the cell populations that preferentially take up these EVs in pulmonary fibrosis. Because pulmonary fibrosis involves multiple cell types that could potentially interact with EVs, defining the predominant recipient population was necessary to guide subsequent functional analyses. We therefore used complementary in vitro and in vivo approaches to compare EV uptake across four representative cell types relevant to pulmonary fibrosis, namely bone marrow-derived macrophages (BMDMs), mouse embryonic fibroblasts (MEFs), MLE-12 pulmonary epithelial cells and mouse microvascular endothelial cells (MMECs).

To identify the predominant recipient population of MSC-derived EVs, we first compared EV uptake across four cell types relevant to pulmonary fibrosis. We incubated equal numbers of DiD-labelled EVs with BMDMs, MEFs, MLE-12 cells, and MMECs for 12 h, then imaged by fluorescence and quantified mean fluorescence intensity (Figure 4K-N). BMDMs showed the highest EV uptake among the four cell types examined, more than 5-fold higher than MLE-12 cells, which showed the second-highest uptake but remained markedly lower than BMDMs. These results identified macrophages as the predominant EV recipient population in vitro system. To determine whether matrix stiffness influenced EV uptake by macrophages, we next compared EVs produced by MSCs cultured on a stiffness-tunable PAA hydrogel platform. EVs derived from MSCs cultured on a stiffness-tunable PAA hydrogel platform showed significantly lower macrophage uptake than those generated on the stiffest matrix (Figure 4J, K), consistent with previous observations that matrix stiffness can modulate the cellular uptake of MSC-derived EVs [10]. To explore the metabolic changes potentially associated with this stiffness-responsive difference in uptake, we further analysed a previously reported stiffness-responsive metabolomic dataset. This analysis revealed enrichment of several cholesterol-related metabolites under the stiff-matrix condition (Figure 4B, C). These findings suggested that stiffness-responsive changes in lipid metabolism, particularly cholesterol-associated pathways, might contribute to the greater macrophage uptake of EVs generated on stiffer matrices [10].

To determine whether the preferential association of MSC-derived EVs with macrophages observed in vitro was also evident in vivo, we administered DiD-labelled EVs to mice 7 days after induction of pulmonary fibrosis and collected lung tissues 48 h later. We then quantified the colocalization of DiD-labelled EV fluorescence with CD68, Vimentin, SFTPC and CD31 to assess EV association with macrophages, fibroblasts, pulmonary epithelial cells and microvascular endothelial cells, respectively. DiD-labelled EV fluorescence showed markedly greater colocalization with CD68⁺ macrophages than with the other three cell populations, with macrophage-associated signals approximately 10-fold higher (Figure 4O, P). These results identified pulmonary macrophages as the predominant EV-associated cell population in pulmonary fibrosis and provided the rationale for focusing subsequent functional analyses on EV-macrophage interactions.

To determine whether chemical activation of ABCA1 altered macrophage uptake of MSC-derived EVs, we compared EVs produced by GSK3987-treated and untreated MSCs. EVs derived from GSK3987-treated MSCs showed greater uptake by macrophages than control EVs (Figure 4J, K), indicating that chemical activation of ABCA1 influenced not only EV production but also their interaction with recipient macrophages. To investigate whether this change in uptake was associated with altered EV membrane composition, we next quantified EV membrane cholesterol. Changes in membrane cholesterol closely paralleled the pattern of macrophage uptake, with EVs containing higher levels of membrane cholesterol showing greater uptake by macrophages (Figure 4H). These findings suggested a link between ABCA1-dependent cholesterol regulation and macrophage uptake of MSC-derived EVs.

Together, these results show that combining a soft matrices condition with chemical activation of ABCA1 increases EV production, elevates EV membrane cholesterol and enhances macrophage uptake. This mechanochemical reprogramming strategy therefore integrates distinct aspects of EV production and delivery that matrix stiffness alone does not simultaneously improve. We therefore examined whether these engineered EVs retained their immunomodulatory activity and whether the enhanced uptake phenotype was attributable to a broader ABCA1-cholesterol dependent mechanism rather than to GSK3987 alone.

### 2.4 Mechanochemical reprogramming coordinates EV production, macrophage uptake and immunomodulatory activity

Previous studies have shown that soft matrices can enhance the immunomodulatory secretory phenotype of MSCs and promote their anti-inflammatory activity [25, 26]. To determine whether the mechanosensitive effect was reproduced in our system, we first assessed MSC-mediated regulation of macrophage polarization using a Transwell co-culture assay. Decreasing matrix stiffness was associated primarily with enhanced M2-like polarization, as indicated by an increased proportion of Arg-1⁺ macrophages. In contrast, the proportion of iNOS⁺ cells varied little under most stiffness-tunable conditions and increased appreciably only at the highest stiffness condition (Figure S4). These results confirmed that soft matrix conditions favoured MSC immunomodulatory activity in the co-culture system.

To determine the specific contribution of MSC-derived EVs to this response, we next isolated EVs from MSCs cultured under stiffness-tunable conditions and treated macrophages with equal numbers of EV particles (Figure 5A), thereby minimizing differences arising from EV abundance. Unlike mechanosensitive pattern observed with MSC co-culture, particle-normalized EV treatment produced a biphasic response. EVs generated at the intermediate stiffness (∼18 kPa) induced the strongest M2-like phenotype, with the highest levels of M2-associated markers and the lowest levels of M1-associated markers, resulting in the highest M2/M1 marker ratio among the groups examined (Figure 5B-D). Thus, the immunomodulatory activity of MSC-derived EVs did not vary monotonically with matrix stiffness, indicating that EV-mediated macrophage regulation reflects mechanosensitive changes in EV properties rather than simply recapitulating MSCs’ overall paracrine response.

**Figure 5.**
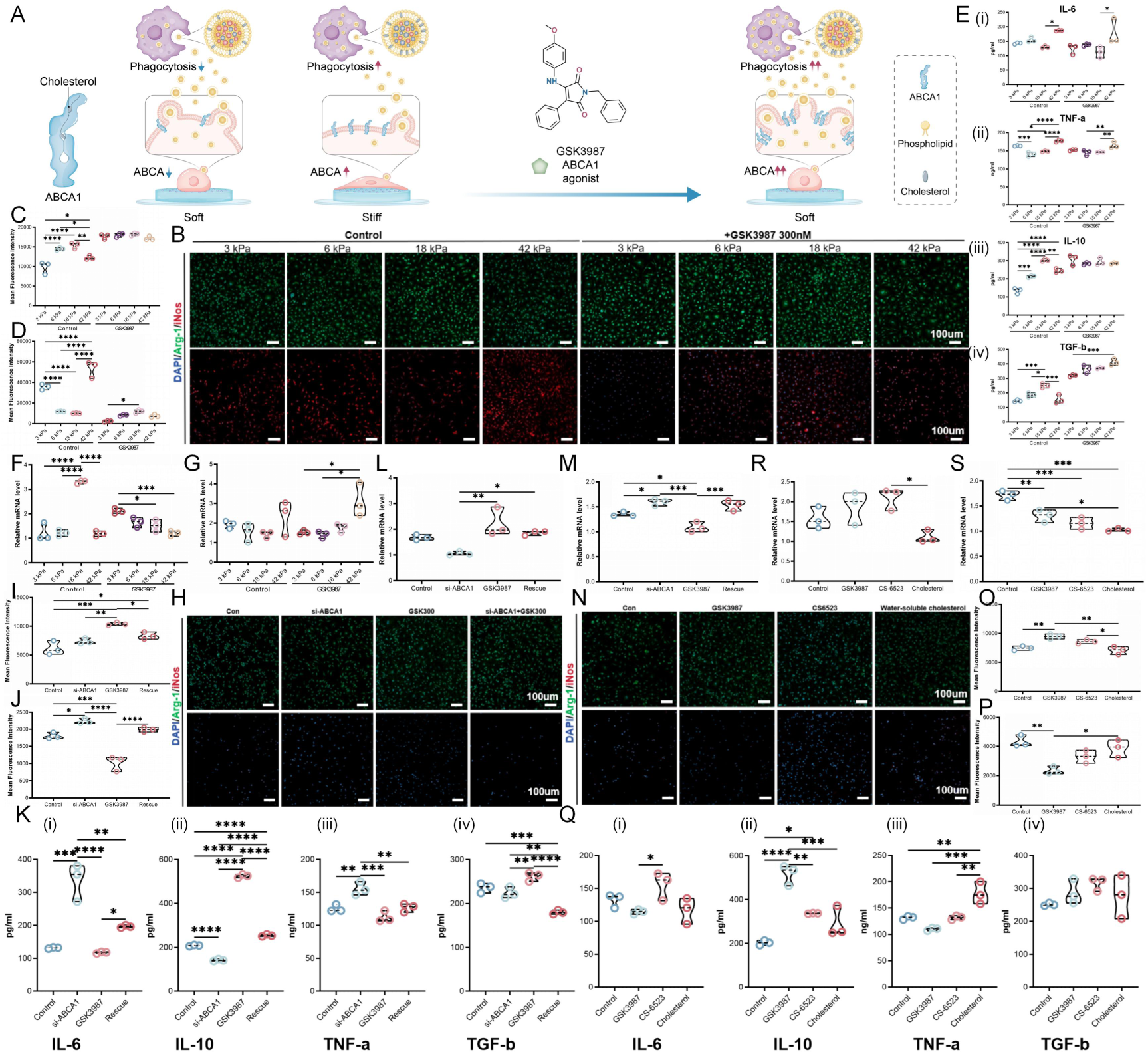
Mechanochemical reprogramming of MSC-derived EV immunomodulatory activity through the ABCA1-cholesterol axis. (A) Schematic of the mechanochemical reprogramming strategy. MSCs cultured on matrices of defined stiffness were chemically modulated for ABCA1 using GSK3987, and the resulting EVs were evaluated for their effects on macrophage polarization. (B) Representative confocal immunofluorescence images of Arg-1 and iNOS in macrophages treated with equal numbers of EV particles derived from MSCs cultured under stiffness-tunable conditions, with or without GSK3987 treatment. (C, D) Quantification of Arg-1 (C) and iNOS (D) mean fluorescence intensity corresponding to (B) (n = 3). (E) ELISA quantification of cytokines in macrophage culture supernatants following treatment with EVs generated under stiffness-tunable conditions and GSK3987 treatment, including (i) IL-6, (ii) TNF-α, (iii) IL-10 and (iv) TGF-β (n = 3). (F, G) RT-qPCR analysis of Arg-1 (F) and iNOS (G) mRNA expression in macrophages treated with the indicated EV preparations (n = 3). (H) Representative confocal immunofluorescence images of Arg-1 and iNOS in macrophages treated with EVs derived from MSCs subjected to si-ABCA1, GSK3987-treated chemical activation of ABCA1, or the indicated rescue treatment. (I, J) Quantification of Arg-1 (I) and iNOS (J) mean fluorescence intensity corresponding to (H) (n = 3). (K) ELISA quantification of cytokines in macrophage culture supernatants following treatment with EVs derived from MSCs subjected to si-ABCA1, GSK3987 treatment, or rescue treatment. (L, M) RT-qPCR analysis of Arg-1 (L) and iNOS (M) expression in macrophages treated with the indicated EV preparations (n = 3). (N) Representative confocal immunofluorescence images of Arg-1 and iNOS in macrophages treated with EVs derived from MSCs subjected to GSK3987, CS6523 or cholesterol treatment. (O, P) Quantification of Arg-1 (O) and iNOS (P) mean fluorescence intensity corresponding to (N) (n = 3). (Q) ELISA quantification of cytokines in macrophage culture supernatants following treatment with EVs derived from MSCs subjected to GSK3987, CS6523 or cholesterol treatment, including (i) IL-6, (ii) TNF-α, (iii) IL-10 and (iv) TGF-β (n = 3). (R, S) RT-qPCR analysis of Arg-1 (R) and iNOS (S) expression in macrophages treated with the indicated EV preparations (n = 3).

To determine whether mechanosensitive changes in EV immunomodulatory activity were associated with molecular cargo, we performed unbiased proteomic profiling of EVs generated under stiffness-tunable conditions. EVs derived from the softest matrix (∼3 kPa) contained a distinct set of upregulated proteins enriched in immune-regulatory pathways, including innate immune activation and inflammatory regulation (Figure S6C). To further assess the immunological relevance of these cargo changes, we compared the proteins enriched in soft-matrix conditioned EVs with an established transcriptomic signature of M2-polarized macrophages (GSE69607) [27]. This analysis identified 36 overlapping targets, including Arginase-1 (Figure S6A, B), indicating that EVs generated under soft-matrix conditions were enriched in proteins associated with an immunoregulatory phenotype.

Overall, these findings suggest that mechanosensitivity influences two important features of MSC-derived EVs. On softer matrices, it promotes an immunoregulatory cargo profile, while on stiffer matrices, it boosts macrophage uptake. This divergence may explain the biphasic immunomodulatory response observed under stiffness-tunable conditions, with EVs generated at intermediate stiffness showing the most favourable combination of cargo characteristics and cellular uptake.

To determine whether chemical activation of ABCA1 could enhance macrophage uptake while preserving the immunomodulatory properties under soft matrix conditions, we treated MSCs with GSK3987 and examined the resulting EVs. GSK3987 increased EV membrane cholesterol (Figure 4H) and enhanced macrophage uptake under stiffness-tunable conditions (Figure 4J, K). Following chemical activation of ABCA1, the differences in macrophage immunomodulatory responses among EVs generated under stiffness-tunable conditions were substantially reduced (Figure 5A). Arg-1 expression no longer differed significantly among the groups (Figure 5B, C), whereas differences in iNOS expression were markedly attenuated compared with those observed before GSK3987 treatment. Notably, macrophages treated with engineered EVs derived from the softest matrix exhibited the lowest iNOS expression, indicating the most pronounced anti-inflammatory phenotype among the conditions examined (Figure 5B, D).

Together, these results support distinct contributions from the mechanical and chemical components of our strategy, with soft matrix conditions favouring an immunoregulatory EV cargo profile and chemical activation of the ABCA1-membrane cholesterol axis enhancing macrophage uptake. Integrating these two components therefore increased EV uptake while retaining the immunomodulatory advantage associated with soft-matrix conditioning, providing the basis for the subsequent mechanochemical reprogramming strategy.

Previous studies have shown that EV membrane composition can influence cellular uptake, with cholesterol enrichment reported to enhance the internalization of milk-derived EVs by tumour cells [28]. Matrix stiffness has likewise been implicated in regulating macrophage uptake of MSC-derived EVs [10]. These observations may explain our previous finding that EVs generated on soft matrices (E ≈ 2 kPa) and stiff matrices (E ≈ 30 kPa) produced comparable therapeutic outcomes in vivo despite differences in their molecular cargo profiles [12]. One possibility is that the greater immunomodulatory potential of soft-matrix conditioned EVs is partly limited by their lower uptake efficiency, thereby reducing the extent to which their functional cargo can be delivered to recipient cells in vivo. This consideration motivated our mechanochemical reprogramming strategy to increase EV membrane cholesterol and improve cellular uptake while preserving the immunomodulatory properties conferred by the soft-matrix condition.

To further validate the effects of matrix stiffness on EV-mediated macrophage regulation beyond immunofluorescence, we assessed macrophage responses at both the transcriptional and cytokine levels. In the absence of GSK3987, RT-qPCR analysis revealed a non-linear mechanosensitive pattern of Arg-1 expression, with significantly higher levels in the intermediate-stiffness groups (Figure 5F). By contrast, iNOS mRNA expression did not differ significantly under stiffness-tunable conditions (Figure 5G). We next quantified secreted inflammatory mediators by ELISA. EVs generated at intermediate stiffnesses generally induced higher levels of the anti-inflammatory cytokines IL-10 and TGF-β, whereas pro-inflammatory cytokine responses varied across readouts, with IL-6 lowest in the 18 kPa group and TNF-α lowest in the 6 kPa group (Figure 5E). These results indicate that no single stiffness produced the most favourable response across all inflammatory markers, although intermediate stiffnesses generally favoured a more anti-inflammatory macrophage phenotype.

To determine whether chemical activation of ABCA1 altered these mechanosensitive functional differences, we next compared macrophage responses to EVs generated from GSK3987-treated MSCs under stiffness-tunable conditions. Following GSK3987 treatment, the differences in Arg-1 and iNOS expression under stiffness-tunable conditions became less pronounced. A similar attenuation was observed at the cytokine level, although the effect was not uniform across all readouts; notably, IL-6 and TNF-α remained higher in the stiffest-matrix condition (Figure 5E). Thus, chemical activation of ABCA1 reduced, but did not completely eliminate, the functional heterogeneity associated with matrix stiffness. Together with the enhanced macrophage uptake observed after GSK3987 treatment, these findings indicate that chemical modulation of the ABCA1-membrane cholesterol axis can improve EV delivery while largely preserving their immunomodulatory activity.

Finally, to identify an EV formulation for subsequent therapeutic evaluation, we considered both immunomodulatory activity and EV production. Although GSK3987-conditioned EVs generated under stiffness-tunable conditions showed broadly similar immunomodulatory profiles, EV production was significantly higher when MSCs were cultured on the softest matrix (∼3 kPa) in combination with GSK3987 (Figure 4G). We therefore selected the 3 kPa/GSK3987 condition for subsequent in vivo studies, as this formulation combined increased EV production with preserved immunomodulatory activity and enhanced macrophage delivery efficiency.

### 2.5 Genetic and chemical validation of the ABCA1-membrane cholesterol axis in MSC-derived EV mediated macrophage immunoregulation

Having established that chemical activation of ABCA1 with GSK3987 under the soft-matrix condition enhanced macrophage uptake while preserving the immunomodulatory activity of MSC-derived EVs, we next sought to determine whether these effects were mediated through the ABCA1-membrane cholesterol axis. To address this question, we used three complementary validation strategies, including (i) genetic perturbation of ABCA1 to assess its functional contribution directly, (ii) chemical activation with a second ABCA1 modulating compound to determine whether the phenotype extended beyond GSK3987, and (iii) direct cholesterol modulation to assess the contribution of membrane cholesterol to the observed EV phenotype.

We first examined the specific contribution of ABCA1 by bidirectionally modulating its expression in MSCs using siRNA-mediated knockdown, chemical activation with GSK3987, and a combined rescue treatment. Si-ABCA1 markedly reduced ABCA1 expression, whereas GSK3987 increased its expression, and combined treatment partially restored ABCA1 levels (Figure S7A-C). MSCs subjected to these perturbations were subsequently cultured on soft matrices (∼3 kPa), and the resulting EVs were collected for downstream characterization and functional analyses (Figure S7G). EVs from these conditions showed comparable size distributions, with particle diameters predominantly ranging from 67 to 79 nm.

To determine whether ABCA1 perturbation also affected EV production, we quantified particle numbers by nanoflow cytometry. EV particle numbers increased to approximately 3-fold above control levels following si-ABCA1, GSK3987, or combined rescue treatment (Figure S7K). Because increased EV production was observed across both loss- and gain-of-function perturbations, these data suggest that experimental modulation of MSCs can influence EV release and that changes in EV yield cannot be attributed solely to the direction of ABCA1 regulation. We therefore focused subsequent analyses on whether changes in ABCA1 expression produced corresponding changes in EV membrane cholesterol and immunomodulatory activity.

We next assessed the effects of these EVs on macrophage polarization. Immunofluorescence analysis showed that EVs generated under GSK3987 treatment induced higher Arg-1 expression, whereas EVs from the si-ABCA1 group were associated with increased iNOS; the rescue group generally displayed an intermediate phenotype (Figure 5H-J). RT-qPCR further showed that increasing ABCA1 activity was associated with higher Arg-1 and lower iNOS expression (Figure 5L, M). Consistent with these findings, ELISA revealed progressive reductions in the pro-inflammatory cytokines IL-6 and TNF-α and increases in IL-10 and TGF-β with increasing ABCA1 activity. However, the magnitude of response varied among individual cytokines (Figure 5K). Together, these results associate ABCA1 activity with a more anti-inflammatory macrophage phenotype induced by MSC-derived EVs, while the intermediate phenotype observed under rescue conditions further supports an ABCA1 dependent component.

To determine whether the effects observed with GSK3987 extended beyond a single compound, we introduced a second ABCA1 agonist, CS-6253, as an independent chemical validation. Dose-response analysis identified 2.5 μg ml^-1^ CS-6253 as the condition associated with the highest ABCA1 expression, and this concentration was used for subsequent experiments (Figure S7D). Western blotting confirmed an approximately 1.5-fold increase in ABCA1 protein following CS-6253 treatment, although the magnitude of activation remained lower than that with GSK3987 treatment (Figure S7E, F). EVs generated under CS-6253 treatment also exhibited a lower zeta potential than those from the control or GSK3987 treatment (Figure S7I), indicating that the two chemical interventions produced distinct changes in EV surface properties.

We next examined whether this independent ABCA1 activation reproduced the increase in EV membrane cholesterol observed with GSK3987. CS-6253 increased EV membrane cholesterol to approximately 4-fold above control levels, although the magnitude of enrichment remained lower than that induced by GSK3987 (Figure S7J). These findings indicate that increased EV membrane cholesterol is reproducible with a second ABCA1-targeting compound and is therefore unlikely to reflect a GSK3987 specific effect. Notably, however, EV particle numbers after CS-6253 treatment remained close to control levels, whereas GSK3987 markedly increased EV yield (Figure S7K). Thus, although both compounds promoted ABCA1-associated enrichment of EV membrane cholesterol, GSK3987 produced a more pronounced increase in EV yield.

To determine whether the two modes of chemical ABCA1 activation produced comparable functional effects, we next assessed macrophage responses to the resulting EVs. EVs generated from GSK3987-treated MSCs induced the most consistent anti-inflammatory phenotype, characterized by increased Arg-1 and reduced iNOS expression (Figure 5N-P). CS-6253-treated EVs produced changes in the same direction, but with a smaller magnitude. RT-qPCR analysis showed a similar pattern (Figure 5R, S), indicating that independent chemical activation of ABCA1 reproduced the overall direction of the GSK3987 induced response, although less strongly.

To determine whether increased membrane cholesterol alone could account for this phenotype, we next examined EVs generated after direct cholesterol treatment. Unlike GSK3987-treated EVs, these EVs did not reproduce the full macrophage polarization response, with macrophages remaining closer to an unpolarized state. Cytokine analysis further distinguished the three treatments. GSK3987-treated EVs reduced IL-6 and TNF-α while increasing IL-10, producing the most consistent anti-inflammatory cytokine profile (Figure 5Q). CS-6253-treated EVs increased IL-10 and TGF-β, but this was accompanied by increased IL-6 relative to control, whereas cytokine levels following direct cholesterol treatment remained broadly comparable to those in the control group.

Together with the genetic perturbation and rescue experiments, these findings support the ABCA1-membrane cholesterol axis as an important contributor to MSC-derived EV mediated macrophage regulation. The partial reproduction of the phenotype by CS-6253 argues against a GSK3987 specific effect. In contrast, the inability of direct cholesterol treatment to fully recapitulate the GSK3987 response indicates that membrane cholesterol contributes to, but does not determine, EV immunomodulatory function. Because GSK3987 provided the most consistent combination of ABCA1 activation, EV membrane cholesterol enrichment, macrophage immunomodulation, and increased EV yield, we selected GSK3987 as the chemical component of the subsequent mechanochemical reprogramming strategy and for in vivo therapeutic evaluation.

### 2.6 Mechanochemically reprogrammed EVs enhance pulmonary retention and abrogate bleomycin-induced fibrosis

To determine whether the improved macrophage uptake observed in vitro translated into altered pulmonary distribution in vivo, we evaluated EV biodistribution in a bleomycin-induced pulmonary fibrosis model following intratracheal administration [29]. Having established that mechanochemically reprogrammed EVs retained immunomodulatory activity in vitro, we compared them with untreated soft-matrix conditioned EVs at equivalent administered doses. To monitor pulmonary distribution over time, we tracked DiD-labelled EVs by longitudinal whole-animal fluorescence imaging. On day 1 after administration, both EV preparations showed widespread pulmonary fluorescence (Figure 6A). The signal subsequently declined in both groups and was no longer readily detectable by day 14 (Figure 6A). Notably, however, mechanochemically reprogrammed EVs maintained higher EV-associated pulmonary fluorescence over the observation period than untreated soft-matrix conditioned EVs (Figure 6B). These findings indicate that mechanochemically reprogrammed EVs were associated with more persistent DiD-labelled EV signals in the lung.

**Figure 6.**
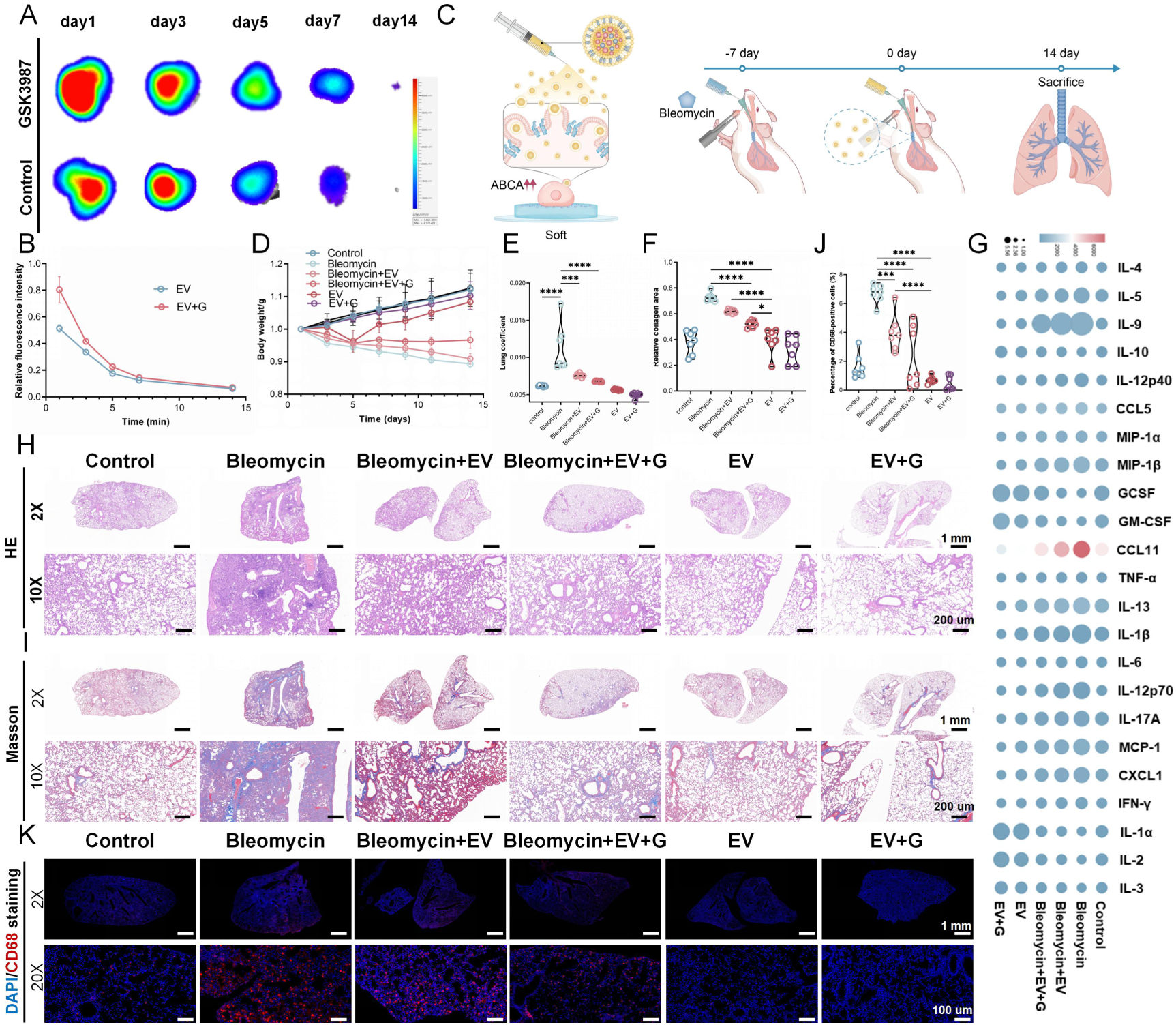
Mechanochemically reprogrammed EVs alleviate pulmonary fibrosis and modulate the pulmonary inflammatory microenvironment. (A, B) Representative longitudinal in vivo fluorescence images of DiD-labelled EVs (A) and quantification of pulmonary DiD-labelled EV fluorescence (B) over 14 days after administration (n = 3). (C) Schematic of the therapeutic study in a bleomycin-induced murine model of pulmonary fibrosis. (D) Longitudinal changes in body weight across treatment groups, expressed relative to the pre-modelling baseline (n = 7). (E) Lung coefficient at day 14 after treatment, calculated as the ratio of lung weight to body weight (n = 7). (F) Quantification of collagen-positive area in lung sections from the indicated treatment groups (n = 7). (G) Multiplex ELISA profiling of cytokines and chemokines in bronchoalveolar lavage fluid (BALF) collected from the indicated treatment groups. (H, I) Representative haematoxylin and eosin (H&E) staining (H) and Masson’s trichrome staining (I) of lung sections at day 14 after treatment, showing lung architecture and fibrotic remodelling. (J) Quantification of CD68⁺ cells in lung sections by immunofluorescence analysis (n = 7). (K) Representative immunofluorescence images of CD68⁺ cells in lung sections at day 14 after treatment, showing macrophage accumulation across treatment groups.

To further assess whether intratracheally administered EVs redistributed to major non-pulmonary organs, we performed ex vivo fluorescence imaging at the corresponding time points. Little or no detectable DiD-labelled EV fluorescence was observed in the major off-target organs examined (Figure S8). Together, these results show that intratracheal delivery largely restricted DiD-labelled EV signals to the lung and that mechanochemically reprogrammed EVs exhibited more persistent pulmonary fluorescence than untreated soft-matrix conditioned EVs, providing a rationale for subsequent evaluation of their therapeutic efficacy in pulmonary fibrosis.

To determine whether the improved delivery properties of mechanochemically reprogrammed EVs translated into greater therapeutic benefit in vivo, we next evaluated disease progression in the bleomycin-induced pulmonary fibrosis model (Figure 6C). Bleomycin-treated mice showed marked body-weight loss, whereas mice receiving either mechanochemically reprogrammed EVs or untreated soft-matrix conditioned EVs resumed weight gain by day 3 after administration. Recovery was more rapid in the mechanochemically reprogrammed EV group than in the untreated soft-matrix conditioned EV group (Figure 6D). Consistent with this improvement, the lung coefficient, an indicator associated with pulmonary oedema and fibrotic burden, was most strongly reduced in mice treated with mechanochemically reprogrammed EVs (Figure 6E). These findings suggest that the enhanced cellular delivery achieved by mechanochemical reprogramming was associated with improved in vivo therapeutic performance.

To further assess the effects of EV treatment on lung pathology, we performed histological analysis 14 days after administration. Bleomycin challenge resulted in extensive alveolar collapse, interstitial thickening, and inflammatory-cell infiltration. In contrast, intratracheal administration of MSC-derived EVs reduced collagen deposition and preserved lung parenchymal architecture, as shown by H&E and Masson’s trichrome staining (Figure 6F, H, I). The protective effect was more pronounced in the mechanochemically reprogrammed EV group, which showed better preserved alveolar structures and less fibrotic remodelling than the untreated soft-matrix conditioned EV group. Immunohistochemical analysis also showed that both EV treatments reduced CD68^+^ macrophage accumulation in the lung, with greater overall improvement after mechanochemical reprogramming (Figure 6K).

To determine whether changes in the pulmonary inflammatory milieu accompanied these histological improvements, we profiled 23 cytokines and chemokines in bronchoalveolar lavage fluid. Mechanochemically reprogrammed EV treatment reduced several pro-inflammatory mediators, including IL-9, IL-6, MIP-1β, CCL11, and IL-1β, while IL-4 and IL-10 remained comparatively stable (Figure 6G). These data indicate that mechanochemically reprogrammed EVs attenuated the pro-inflammatory lung microenvironment, alongside improvements in tissue pathology.

Together, these results show that intratracheal administration of mechanochemically reprogrammed EVs produced greater overall therapeutic benefit than untreated soft-matrix conditioned EVs, as reflected by faster body-weight recovery, a lower lung coefficient, reduced fibrotic remodelling and a less inflammatory pulmonary microenvironment. The concordance between improved pulmonary delivery and therapeutic outcome further supports using mechanochemical reprogramming to enhance the in vivo performance of MSC-derived EVs.

### 2.7 Multi-omics integration and machine learning identify GLOD4 as the key molecular effector mediating immunomodulatory capacity of engineered EVs

To identify EV-associated molecular candidates relevant to human pulmonary fibrosis, we integrated nine independent transcriptomic datasets from patients with idiopathic pulmonary fibrosis (IPF) and non-fibrotic controls (Figure 7A). To improve comparability across datasets, expression data were normalized and corrected for inter-study batch effects before samples were stratified into IPF and control groups (Figure 7B) [30]. To prioritize EV cargo relevant to the disease-associated molecular changes observed in human IPF, we next integrated the clinical transcriptomic data with the proteomic profile of immunomodulatory EVs generated under soft-matrix conditions. Specifically, proteins enriched in soft-matrix conditioned EVs were intersected with genes downregulated in human IPF tissues. This cross-omics analysis identified 12 overlapping candidates that were reduced in IPF while being enriched in the EV proteome (Figure 7C), providing a focused set of molecules for subsequent prioritization.

**Figure 7.**
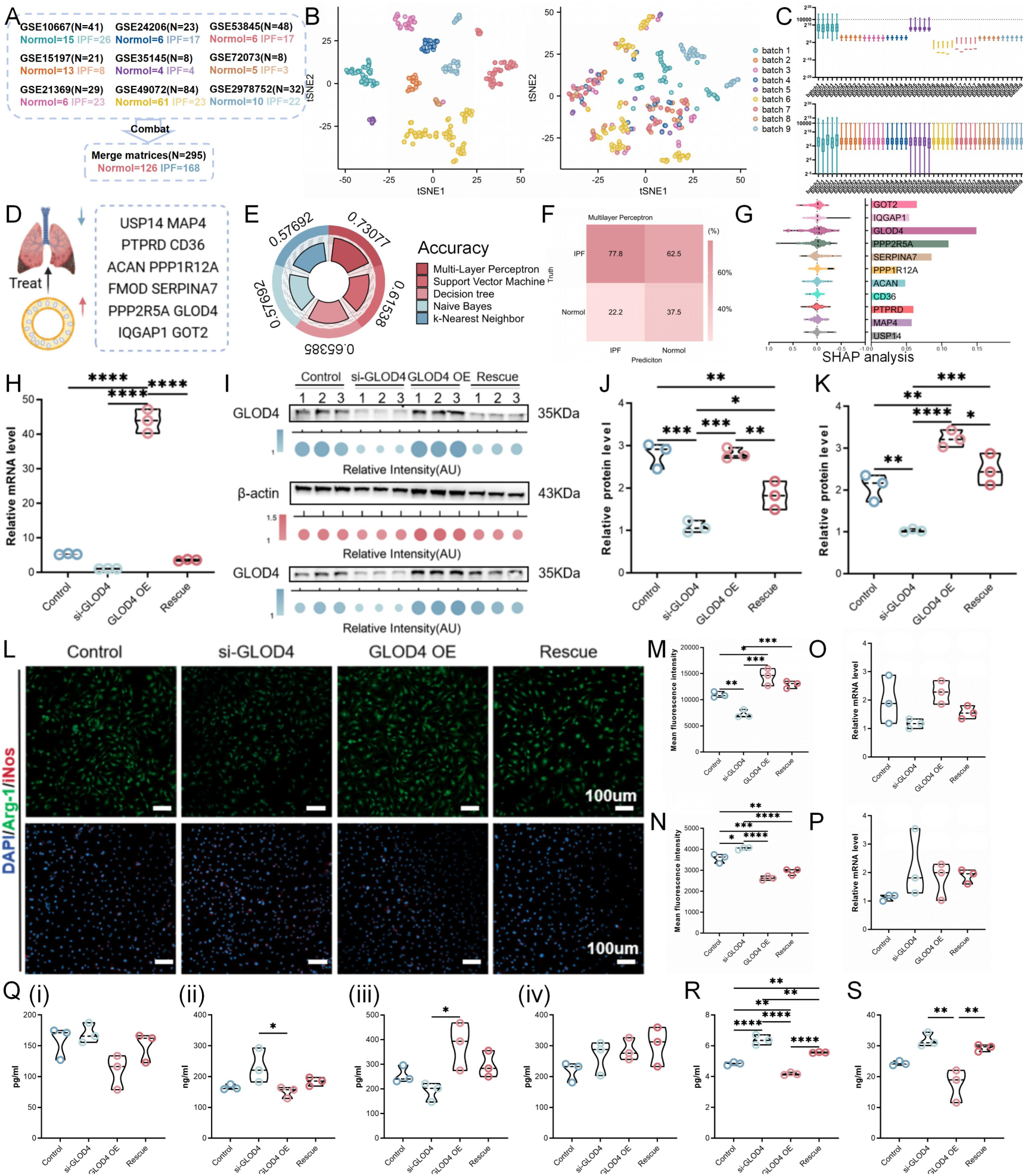
Machine learning-guided identification and functional validation of GLOD4 as an immunomodulatory cargo of MSC-derived EVs in pulmonary fibrosis. (A) Summary and statistical characterization of publicly available human pulmonary fibrosis transcriptomic datasets used for integrative analysis. (B) t-SNE visualization of sample distributions before and after ComBat correction, showing reduced batch-associated variation. (C) Quantitative comparison of batch distributions before and after ComBat correction. (D) Identification of candidate therapeutic EV proteins by integrating public pulmonary fibrosis datasets with our MSC-derived EV proteomic dataset. (E) Classification performance of five machine-learning algorithms in distinguishing individuals with pulmonary fibrosis from healthy controls using the selected EV-associated protein features. (F) Confusion matrix showing the classification performance of the optimized multilayer perceptron (MLP) model. (G) SHAP analysis showing the relative contribution of the selected EV-associated protein features to model prediction and identifying GLOD4 as a leading contributor. (H) RT-qPCR analysis of GLOD4 mRNA expression in MSCs following siRNA-mediated GLOD4 knockdown (si-GLOD4), GLOD4 overexpression (GLOD4 OE), or rescue (Rescue) treatment (n = 3). (I) Western blot analysis of GLOD4 protein expression in MSCs following si-GLOD4, GLOD4 OE, or Rescue treatment. (J) Quantification of GLOD4 protein expression in MSCs corresponding to (I) (n = 3). (K) Quantification of GLOD4 protein abundance in EVs secreted by MSCs subjected to si-GLOD4, GLOD4 OE, or Rescue treatment (n = 3). (L) Representative immunofluorescence images of the macrophage polarization markers Arg-1 and iNOS after treatment with equal numbers of EV particles derived from MSCs subjected to si-GLOD4, GLOD4 OE, or Rescue treatment. (M, N) Quantification of the mean fluorescence intensity of Arg-1 (M) and iNOS (N) corresponding to (L) (n = 3). (O, P) RT-qPCR analysis of Arg-1 (O) and iNOS (P) expression in macrophages treated with equal numbers of the indicated MSC-derived EV particles (n = 3). (Q) ELISA quantification of cytokines released into macrophage culture supernatants following treatment with the indicated EV preparations: IL-6 (i), IL-10 (ii), TNF-α (iii), and TGF-β (iv) (n = 3). (R, S) Quantification of the GLOD4-associated downstream products methylglyoxal (MGO) (R) and Nε-(1-carboxyethyl)lysine (CEL) (S) in macrophages following treatment with EVs derived from MSCs subjected to si-GLOD4, GLOD4 OE, or Rescue treatment (n = 3).

To further refine these candidates according to their ability to distinguish IPF from control samples, we applied recursive feature elimination with cross-validation (RFECV). FMOD contributed least to classification performance and was removed, leaving 11 candidate features for subsequent modelling (Figure 7D). We then compared five machine-learning classifiers trained using these candidates. Among the models examined, the multilayer perceptron (MLP) achieved the highest classification performance, with 73.07% accuracy in distinguishing IPF from control samples (Figure 7E, F). To determine which retained candidates contributed most to model predictions, we next interpreted the MLP classifier using SHapley Additive exPlanations (SHAP). GLOD4 showed the highest feature contribution to discrimination between IPF and non-fibrotic samples (Figure 7G). Notably, GLOD4 was reduced in the human IPF datasets while being enriched in the immunomodulatory EV proteome. Together, these independent lines of evidence led us to prioritize GLOD4 as a candidate functional EV cargo for subsequent experimental validation of its contribution to macrophage immunoregulation and the therapeutic activity of soft-matrix conditioned EVs.

The glyoxalase gene family comprises a critical regulatory cluster of six enzymes characterized by distinct structural and functional properties, including GLOD4 [31]. These family members orchestrate diverse physiological processes spanning amino acid metabolism exemplified by 4-hydroxyphenylpyruvate dioxygenase (HPD), primary metabolism mediated by methylmalonyl-CoA epimerase (MCEE), and aldehyde detoxification governed by glyoxalase 1 (GLO1). Consequently, the perturbation of these enzymatic networks is collectively associated with significant disease outcome [32]. The canonical function of this enzymatic family is the detoxification of reactive dicarbonyls that would otherwise react with cellular nucleophiles to precipitate deleterious modifications of lipids proteins and DNA. These destructive modifications subsequently activate canonical stress responses, including the heat shock response, unfolded protein response, and DNA damage response. Previous studies have confirmed that glyoxalase family proteins can modulate pulmonary fibrosis and other pathologies mediated by advanced glycation end products [33]. Consequently, glyoxalases are essential for maintaining cellular homeostasis and preventing the progression of metabolic diseases. Specifically, GLOD4 has been identified in extracellular EVs and mitochondria, which corroborates the exosomal proteomics findings of this study [34].

To determine whether GLOD4 contributes functionally to the immunomodulatory activity of MSC-derived EVs, we first established a donor-cell genetic manipulation system incorporating siRNA-mediated GLOD4 knockdown (si-GLOD4), GLOD4 overexpression (GLOD4 OE), or rescue (Rescue) treatment. RT-qPCR and Western blotting confirmed the expected changes in GLOD4 expression at both the mRNA and protein levels in MSCs (Figure 7H-J). We then examined GLOD4 abundance in EVs released from these genetically manipulated MSCs. Western blotting showed that EV-associated GLOD4 changed in the same direction as GLOD4 expression in the corresponding donor cells (Figure 7J, K), establishing an experimental system to selectively modulate GLOD4 cargo in MSC-derived EVs.

To determine how changes in EV-associated GLOD4 affected macrophage immunophenotypes, we treated macrophages with the corresponding EV preparations and assessed Arg-1 and iNOS expression by immunofluorescence. Increasing EV-associated GLOD4 was associated with higher Arg-1 and lower iNOS expression, whereas GLOD4 knockdown produced the opposite trend and the rescue condition partially restored the phenotype (Figure 7L-N). This pattern was consistent with the anti-inflammatory phenotype induced by soft-matrix conditioned EVs and supports a functional contribution of EV-associated GLOD4 to macrophage immunoregulation.

To determine whether these phenotypic changes were accompanied by broader transcriptional polarization, we next quantified Arg-1 and iNOS mRNA expression by RT-qPCR. Different from the immunofluorescence findings, neither transcript differed significantly among the GLOD4 manipulated EV groups, indicating that modulation of EV-associated GLOD4 alone had only a limited effect on the transcriptional polarization programme (Figure 7O, P). We further examined secreted cytokines to determine whether GLOD4 preferentially affected specific components of the macrophage inflammatory response. Increasing EV-associated GLOD4 was accompanied by increased IL-10 and reduced TNF-α secretion, whereas IL-6 and TGF- β remained largely unchanged across the groups (Figure 7Q). Together, these findings identify GLOD4 as a functionally relevant component of the immunomodulatory cargo of MSC-derived EVs, while indicating that its effects are selective and insufficient to account for the full macrophage response elicited by soft-matrix conditioned EVs.

To determine the extent to which GLOD4 alone could account for the immunomodulatory effects of mechanochemically reprogrammed EVs, we compared the magnitude of the responses induced by GLOD4-manipulated EVs with those observed following ABCA1-membrane cholesterol-based mechanochemical reprogramming. GLOD4 manipulation produced substantially smaller changes, indicating that although EV-associated GLOD4 contributes to macrophage immunoregulation, modulating this single cargo is insufficient to reproduce the broader functional phenotype of the engineered EVs. These findings suggest that the therapeutic activity of MSC-derived EVs reflects the combined effects of membrane composition, recipient-cell uptake and multiple functional cargos rather than the action of a single molecular determinant.

To determine whether EV-associated GLOD4 engaged its expected downstream molecular programme in recipient macrophages, we next quantified the GLOD4-associated molecules methylglyoxal (MGO) and Nε-(1-carboxyethyl)lysine (CEL). Both showed an inverse relationship with EV-associated GLOD4 abundance, with their levels decreasing as EV GLOD4 increased. These results indicate that genetic modulation of GLOD4 in donor MSCs is transmitted through the EV cargo and is sufficient to alter GLOD4-associated downstream responses in recipient macrophages.

Together, these genetic perturbation studies establish GLOD4 as a functionally relevant component of the immunomodulatory cargo of soft-matrix conditioned MSC EVs. Increasing EV-associated GLOD4 promoted several anti-inflammatory features, including higher Arg-1 and IL-10 and lower TNF-α, but did not fully reproduce the immunomodulatory phenotype induced by mechanochemically reprogrammed EVs. GLOD4 therefore contributes to, but does not solely determine, the therapeutic activity of MSC-derived EVs.

## 3 Discussion

Contemporary efforts in biomimetic EV engineering, whether building soft matrices that recapitulate physiological microenvironments or developing functionalized microcarriers, fundamentally face an intrinsic mechanics-induced functional paradox [10, 12]. Specifically, soft matrices promote multivesicular body fusion and release by alleviating cytoskeletal tension, thereby increasing EV secretion yield and preserving superior immunomodulatory activity [35]. However, soft matrix conditions inadvertently dampen cellular lipid metabolic networks, leading to cholesterol-deficient EV membranes that severely impair their internalization by recipient cells. Conversely, stiff matrices increase EV membrane cholesterol through ABCA1 upregulation, but at the cost of reduced EV yield and a pro-inflammatory cargo profile, compromising immunomodulatory activity. Furthermore, although fluid shear stress or energy-wave stimulation can drastically increase EV yield, excessive physical stimulation often risks compromising cellular viability and exacerbating cargo heterogeneity [36, 37]. This biological dichotomy within physical cue transduction pathways indicates that relying exclusively on uniaxial physical microenvironmental mimicry has reached a translational bottleneck, making it difficult to concurrently satisfy clinical demands for high yield, high immunomodulatory activity, and high delivery efficiency with matrix stiffness as the sole regulatory cue. Consequently, multidimensional regulatory strategies are urgently required to reach the optimal treatment solution.

To overcome this limitation, this study moves beyond traditional empirical parameter optimization by using bioinformatic analysis as a mechanistic compass. Through unbiased transcriptomic screening, we precisely pinpointed ABCA1 as the pivotal mechanosensor linking extracellular mechanical signals to intracellular lipid metabolism. This discovery reveals that the molecular etiology of the deficient delivery efficiency observed in soft-matrix conditioned EVs lies in specific reprogramming of cholesterol metabolism, providing a solid theoretical underpinning for our proposed mechanochemically reprogrammed strategy. Briefly, this strategy exploits the physical microenvironment of soft matrices to preserve EV immunomodulatory activity. Meanwhile, chemical activation of the ABCA1 pathway increases membrane cholesterol levels, promoting macrophage phagocytosis and boosting EV production. As a result, these mechanochemically reprogrammed EVs exhibit high yield, strong immunomodulatory capacity, and high delivery efficiency, offering a novel paradigm for overcoming clinical translation bottlenecks in nanomedicine.

Aligning with recent research demonstrating that hemodynamic forces from hypertension upregulate ABCA1 expression in endothelial cells to promote lipid efflux [38] and that matrix stiffness dictates neutral lipid excretion in stem cells [23], our results confirm that matrix stiffness positively regulates ABCA1 expression and acts as a mechanical switch for lipid metabolic reprogramming. Optimizing membrane cholesterol content has emerged as a critical strategy to potentiate EV delivery, as representative studies highlight that membrane composition governs intracellular bioavailability and lysosomal escape [28, 39]. While cargo defines therapeutic potential, lipid structure dictates successful cellular entry. Current cholesterol-modification methods, which rely heavily on synthetic nanoparticles, often lack the bioactivity and immunocompatibility inherent to cell-derived EVs [37]. We bridge this gap by harnessing the mechanosensitive-ABCA1 axis to engineer native EVs. This approach successfully generates engineered EVs characterized by robust secretion, superior delivery efficiency, and potent therapeutic efficacy.

## 4 Conclusion

In summary, we have unveiled that increasing matrix stiffness upregulates ABCA1 expression, which in turn regulates exosomal lipid composition and internalization efficiency. Building on this mechanism, we devised a mechanochemically reprogrammed strategy that integrates soft-matrix conditions with chemical activation to overcome the inherent trade-off in natural EVs, characterized by a dichotomy between cargo potency and delivery efficiency. This mechanochemically reprogrammed approach yielded engineered EVs with robust pulmonary retention and potent immunomodulatory activity. Moreover, advanced machine learning analysis of clinical datasets pinpointed GLOD4 as the indispensable therapeutic payload enriched within soft-matrix conditioned EVs responsible for abrogating macrophage inflammation. Collectively, our study presents a paradigm shift from passive isolation to rational EV programming, and offers a potent non-invasive therapeutic modality for a broad spectrum of fibrotic pathologies that reconciles biological activity and target cell uptake (Figure 8).

**Figure 8.**
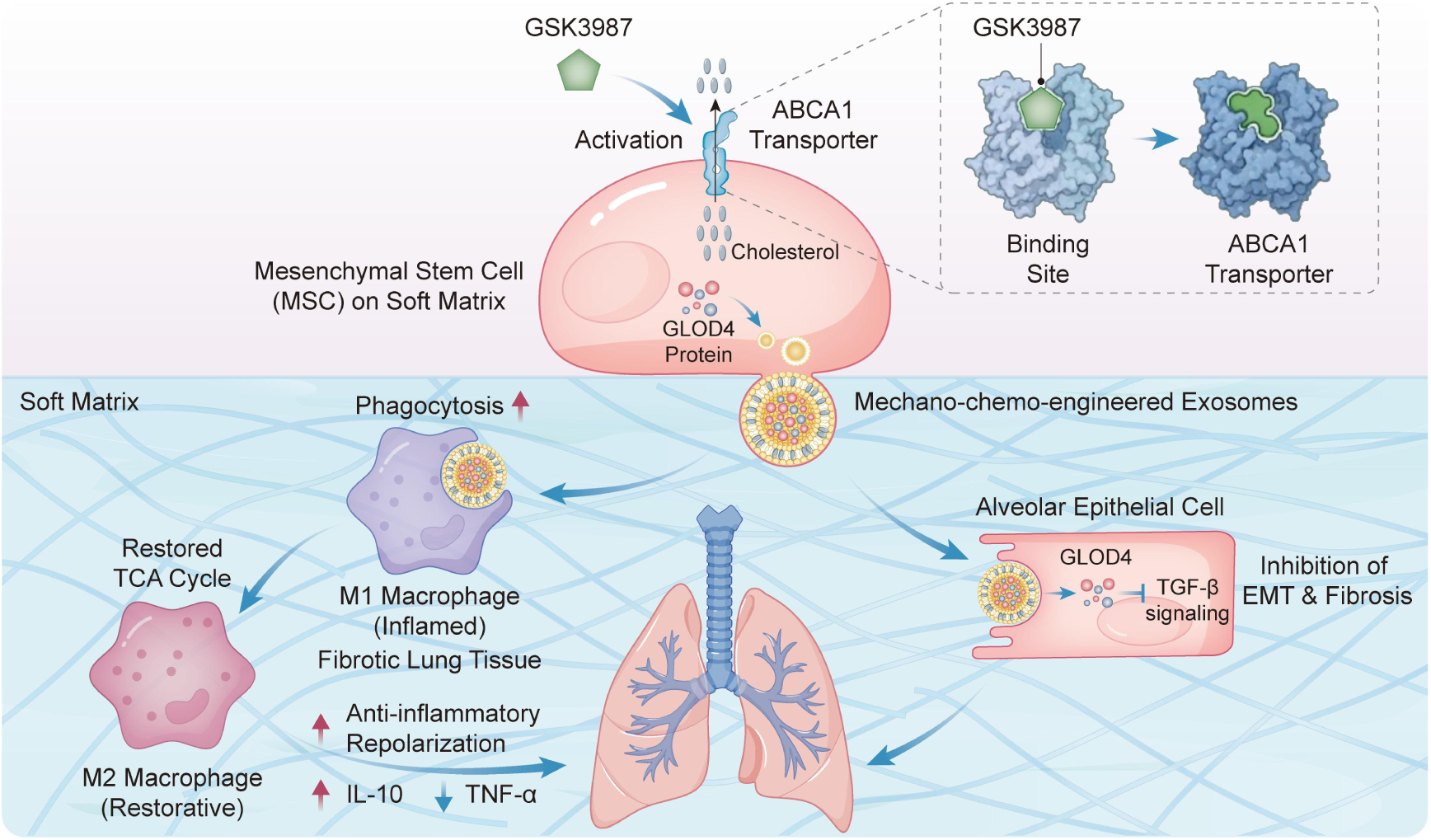
A mechanochemically reprogrammed strategy to engineer MSC-derived EVs with enhanced immunomodulatory efficacy.

## 5 Experimental Section

### 5.1 Data analysis

#### 5.1.1 Bioinformatics analysis

We initially searched publicly available human pulmonary fibrosis transcriptomic datasets and identified nine relevant datasets, including GSE10667 (n = 41), GSE24206 (n = 23), GSE53845 (n = 48), GSE15197 (n = 21), GSE35145 (n = 8), GSE72073 (n = 8), GSE21369 (n = 29), GSE49072 (n = 84) and GSE2978752 (n = 32) included in the integrated analysis. After integration, the combined dataset comprised 126 control samples and 168 pulmonary fibrosis samples. Each sample represented an independent biological specimen.

Limma (linear models for microarray data, DOI:10.1093/nar/gkv007) is a differential expression screening method based on the generalized linear model. In this study, the R package limma (version 3.40.6) was used for differential expression analysis to identify genes differentially expressed between the soft and stiff matrix groups. KEGG Pathway and GO enrichment analyses were performed in R 4.2.1, utilizing the following R packages: clusterProfiler (for enrichment analysis) and org. Hs.eg.db (for ID conversion), and GOplot (for z-score calculation). The specific processing procedure was as follows: after ID conversion of the input molecule list, enrichment analysis was conducted using the clusterProfiler package; the z-score values corresponding to each enriched term were calculated via the GOplot package using the provided molecular values; and further analysis was performed in combination with Cytoscape (version 3.9.1) and the ClueGO plugin (version 2.5.9) [40].

#### 5.1.2 Machine learning

We corrected batch-associated variation across datasets using ComBat. To identify EV-associated candidate therapeutic features, we focused on molecules that were increased in EVs generated from MSCs cultured on the 3 kPa soft matrix and decreased in pulmonary fibrosis samples. The intersection of these two datasets yielded 12 candidate molecules. Then these candidates were subjected to recursive feature elimination with cross-validation (RFECV), retaining 11 features for machine-learning analysis. The integrated cohort was divided into training, validation, and test subsets at a 7:2:1 ratio, assigning each biological sample to only one subset.

REFCV is an advanced optimization of the Recursive Feature Elimination (RFE) method that integrates cross-validation to provide a more rigorous feature-selection framework [41]. This approach systematically removes features that contribute less to the model, gradually reducing feature-space dimensionality and avoiding interference from redundant information and irrelevant variables. In this process, cross-validation serves as a critical evaluation step. It runs multiple rounds of training and validation across different feature subsets, using model performance metrics as quantitative criteria to select the feature combination that maximizes the model’s generalization.

Classification evaluation metrics are primarily derived by comparing ground-truth and predicted labels. Specifically, True Positives (TP) denote the number of correctly classified positive instances, True Negatives (TN) represent the number of correctly classified negative instances, False Positives (FP) refer to the number of incorrectly classified positive instances (i.e., instances actually belonging to the negative class but predicted as positive), and False Negatives (FN) indicate the number of incorrectly classified negative instances (i.e., instances actually belonging to the positive class but predicted as negative).

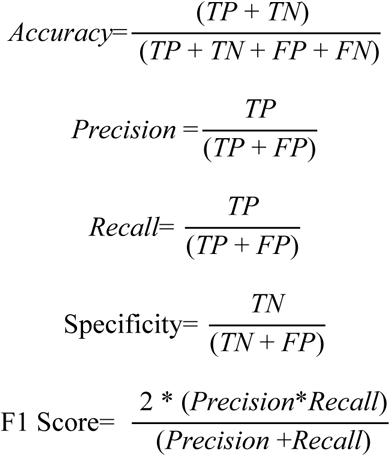

The closer the aforementioned classification metrics are to 1, the better the results.

k-Nearest Neighbors (k-NN) is a fundamental classification algorithm that offers advantages such as ease of use, no training required, and robustness to outliers [42]. In this study, the model was implemented by calling the built-in fitcknn function in MATLAB. Naive Bayes classification is a classification algorithm based on Bayes’ theorem, whose core assumption is that features are mutually independent, i.e., each feature contributes independently to the classification result [43]. This assumption simplifies conditional probability calculations, allowing the overall conditional probability to be estimated from each feature’s probability independently. The model was implemented by calling MATLAB’s built-in fitcnb function. Decision tree classification is a tree-structure-based classification algorithm that offers several advantages, including ease of understanding and interpretation, the ability to handle both discrete and continuous features, and insensitivity to outliers [44]. In this study, the model was implemented by calling the built-in fitctree function in MATLAB. The Support Vector Machine (SVM) is a classic classification algorithm that offers advantages such as high accuracy, the ability to handle high-dimensional data, and good performance on small-sample datasets [45]. However, it also has limitations, including limited ability to handle large-scale datasets and multi-class classification problems, as well as sensitivity to noise and outliers. In this study, the model’s classification function was implemented by calling the built-in fitcecoc function in MATLAB. As a typical feedforward neural network model, the Multilayer Perceptron (MLP) is essentially a multi-layer nonlinear mapping system composed of several hidden layers and input/output layers [46]. Its basic unit is an artificial neuron that simulates biological neurons, enabling layer-by-layer abstraction and high-order feature extraction through weighted connections and activation functions. In MATLAB, the fitrnet function in the Neural Network Toolbox can be used to construct the MLP model.

#### 5.1.3 Statistical analysis

For mouse body weight and fluorescence intensity measured at consecutive time points, the data are presented as mean ± standard deviation (mean ± SD). Statistical analyses were performed using GraphPad Prism. Normality and homogeneity of variance were assessed before statistical testing, and homogeneity of variance among groups was evaluated using the Brown-Forsythe and Bartlett tests. One-way ANOVA was used for overall comparisons among treatment groups.

For datasets presented as violin plots, all individual data points are shown. After confirming normality and homogeneity of variance, overall comparisons among multiple groups were performed using one-way ANOVA, followed by Tukey’s post hoc multiple-comparisons test for pairwise comparisons. P < 0.05 was considered statistically significant.

In heatmap-based visualizations, each circle represents the mean value for the corresponding experimental group. In heatmaps of bronchoalveolar lavage fluid cytokine profiles, the color gradient represents the absolute expression level of each cytokine, while circle size reflects the cytokine’s normalized value and facilitates visual comparison of relative differences among groups.

### 5.2 Hydrogel fabrication

#### 5.2.1 Hydrogel fabrication

Soft or stiff matrices were fabricated by maintaining the acrylamide ( 99%, CAS No. 79-06-1, Cat. No. A15120-1kg, Molecular formula: C^3^H_5_NO, MW: 71.08) was purchased from Shanghai Acmec Biochemical Co., Ltd.) concentration constant (8% (w/v)) and N-acryloxysuccinimide (NAS; ≥97%, CAS 38862-24-7, C_7_H_7_NO_4_, MW 169.13; Bidepharm, Shanghai, China; Cat. No. BD4868) (0.8% (w/v)), while varying the concentration of bisacrylamide (99%, CAS 110-26-9, C_7_H_10_N_2_O_2_, MW 154.17; Macklin Biochemical, Shanghai, China; Cat. No. N917391) (0.05-0.8% (w/v)). To initiate photocrosslinking, photoinitiator 2-Hydroxy-1-[4-(2-hydroxyethoxy)phenyl]-2-methyl-1-propanone (Irgacure 2959, I2959; 98%, CAS 106797-53-9, C_12_H_17_O_4_, MW 224.25; MilliporeSigma, Cat. No. 410896) (0.5% (w/v)) was used. Subsequently, the hydrogel surfaces were incubated overnight at 4°C in 0.2mg/ml type I collagen (rat tail, Corning CB40236). Uncoated collagen and residual acrylamide monomers were removed by PBS washes, and unreacted N-acryloxysuccinimide groups were blocked with serum-free medium to achieve swelling equilibrium.

#### 5.2.2 Mechanical test

Prior to stress-strain measurements, 800 μL of prepolymer solution was added to cylindrical plastic molds (diameter: 12 mm, height: 8 mm) in accordance with ISO 7743 standards, followed by UV curing to crosslink. The compressive properties of polyacrylamide hydrogels were tested using a mechanical testing machine (E43, MTS Instruments, USA) at humidity above 60%. The elastic modulus was calculated based on the initial slope of the linear region (2%-8% strain) of the stress-strain curve.

#### 5.2.3 Rheological measurements

Rheological measurements were conducted using a hybrid rheometer (DHA, TA Instruments, USA). Samples were tested at 25 °C, with all measurements performed on a 20 mm flat steel plate geometry. Oscillatory shear tests were carried out at a constant frequency (1 Hz) and variable shear strain (γ) to determine the storage modulus (G’) and loss modulus (G") across a frequency range of 0.1-100 rad/s.

#### 5.2.4 Scanning Electron Microscopy

Polyacrylamide hydrogels were first frozen at -80°C and then freeze-dried to remove moisture. To enhance their electrical conductivity, the hydrogels were sputtered with gold. The pore sizes of hydrogels with different compositions were observed using a scanning electron microscope (SEM) and quantified using ImageJ software.

#### 5.2.5 Contact angle measurement

The contact angles of polyacrylamide hydrogels with different ratios were measured using the sessile drop method: a 10 μL deionized water droplet was placed on the hydrogel surface, and a contact angle goniometer automatically measured the contact angle.

### 5.3 Cell culture

#### 5.3.1 Stem cell culture

Human umbilical cord MSCs were generously provided by Dr. He from the Second Affiliated Hospital of Dalian Medical University (KY2025-186-01). The cells were cultured at 37 °C in a humidified atmosphere containing 5 % CO2 in the growth medium containing α-MEM (α-Minimum Essential Medium), 10 % fetal bovine serum (FBS), 1% penicillin/streptomycin (all reagents purchased from Thermo Fisher, Shanghai, China), and were passaged at 70-80 % confluency and used at passage 5.

Immortalized human bone marrow-derived mesenchymal stem cells (HMSC-bm-Immortalized; cat. no. ZQY003; Shanghai Zhong Qiao Xin Zhou Biotechnology Co., Ltd.) were cultured in the corresponding complete medium for immortalized cells (cat. no. ZMY003) at 37 °C in a humidified incubator with 5% CO_2_. The culture medium was replaced every 2-3 days. When cells reached approximately 80% confluence, they were detached using 0.05% trypsin (cat. no. CSP048) and passaged at a 1:2 ratio. Cells from passage 10 were used for all experiments.

#### 5.3.2 Macrophage culture

Mouse monocytes were extracted from the bone marrow of 4-week-old male C57/BL6 mice. Briefly, mononuclear cells were separated and cultured in α-MEM containing 10 % FBS, 2 mM L-glutamine, 100 U/ml penicillin/streptomycin, and 30 ng/mL Macrophage Colony-Stimulating Factor (M-CSF, PeproTech, 315-02) for 24 h. Non-adherent cells were cultured in an incubator at 37 °C and 5% CO2 with M-CSF for an additional 3 days, until they reached 80% confluency. LPS (100 ng/mL, Solarbio, China; IL2020) and IFN-γ (20 ng/mL, Peprotech, 315-05) were co-incubated with macrophages for 24 h at 37 °C to induce the proinflammatory phenotype. After induction, the supernatant was collected, and the proinflammatory macrophages were washed three times with PBS to remove the inducer and other components (DUTSBE250827-02).

#### 5.3.3 Mouse fibroblast culture

Mouse fibroblasts (Shanghai Zhong Qiao Xin Zhou Biotechnology Co., Ltd.) were maintained as adherent cultures at 37 °C in a humidified incubator with 5% CO2. Cells were preferably cultured in the corresponding complete medium supplied by the manufacturer; alternatively, a conventional formulation consisting of high-glucose DMEM supplemented with 10% fetal bovine serum, 1% penicillin-streptomycin, GlutaMAX, and non-essential amino acids was used (cat. no. ZQ-104). The culture medium was replaced every 2-3 days. Cells were passaged at approximately 80-85% confluence at a routine split ratio of 1:2.

#### 5.3.4 Mouse microvascular endothelial cell culture

Mouse microvascular endothelial cells (bEnd.3; Shanghai Zhong Qiao Xin Zhou Biotechnology Co., Ltd.) were cultured as adherent cells at 37 °C in a humidified atmosphere containing 5% CO2. Cells were maintained in the corresponding complete medium (cat. no. ZM0090), based on high-glucose DMEM supplemented with 10% fetal bovine serum and 1% penicillin-streptomycin. To promote cell attachment, we precoated culture vessels with 0.1% gelatin at 37 °C for 1 h before seeding cells. The culture medium was replaced every 2-3 days, and cells were passaged at approximately 80% confluence at a split ratio of 1:3 to 1:4.

#### 5.3.5 MLE-12 mouse alveolar epithelial cells culture

MLE-12 mouse alveolar epithelial cells were maintained as adherent cultures at 37 °C in a humidified incubator with 5% CO2. Cells were cultured in HITES medium based on DMEM/F12 supplemented with 2% fetal bovine serum, 1% penicillin-streptomycin, L-glutamine, insulin, transferrin, sodium selenite, hydrocortisone, and β-estradiol. The culture medium was replaced every 2-3 days. Cells were passaged at approximately 80% confluence at a split ratio of 1:2 to 1:3.

### 5.4 Preparation and characterization of EVs

#### 5.4.1 Isolation of EVs

MSCs were seeded onto polyacrylamide hydrogels of defined stiffness and cultured in complete medium until they attached. The cells were then washed three times with PBS to remove residual serum-containing medium, then replaced with serum-free α-MEM. After a further 48 h of culture, conditioned medium was collected for extracellular vesicle (EV) isolation.

The conditioned medium was first filtered through a 0.22-μm membrane to remove residual hydrogel fragments and large particulate contaminants, then centrifuged at 5,000 rpm to remove remaining cell debris and apoptotic bodies. The resulting supernatant was processed using either polyethylene glycol (PEG)-based precipitation or ultrafiltration, depending on the downstream application.

For PEG-based EV enrichment, add PEG 8000 (Sigma-Aldrich, CAS no. 25322-68-3) to the clarified conditioned medium to a final concentration of 8% (w/v). After thorough mixing, samples were incubated overnight at 4 °C and centrifuged at 10,000 rpm for 1 h. The resulting EV-containing pellet was resuspended in PBS at one-fifth of the original conditioned-medium volume. PEG 8000 was then added to a final concentration of 5% (w/v), followed by incubation at 4 °C for at least 4 h and centrifugation at 10,000 rpm for 45 min. The resulting pellet was resuspended in PBS and used primarily for Western blot analysis of EV-associated protein markers.

For ultrafiltration-based EV enrichment, 100 kDa molecular-weight-cut-off Amicon Ultra centrifugal filters were pre-equilibrated with sterile PBS before use. Clarified conditioned medium was loaded onto the filtration units and centrifuged at 4 °C and 4,000 × g using a swinging-bucket rotor until the retentate volume was reduced to approximately 400-500 μl. Sterile PBS was then added to the retentate, and the ultrafiltration step was repeated one to two times to achieve buffer exchange and reduce soluble protein contaminants. The EV-enriched retentate was gently recovered from the membrane surface and used for nanoflow cytometry, transmission electron microscopy, and subsequent functional assays.

#### 5.4.2 Phagocytosis assay of EVs

EVs (1 mg ml⁻¹) were incubated with DiD fluorescent dye (5 μM; Thermo Fisher Scientific, V22887) at 37 °C for 20 min in the dark. To remove unincorporated dye, add PEG 8000 to the EV suspension to a final concentration of 5% (w/v), incubate at 4 °C for at least 4 h, and centrifuge at 10,000 rpm for 45 min. Discard the supernatant, then resuspend the EV-containing pellet in PBS. This washing procedure was repeated three times to minimize residual free DiD.

For cellular uptake assays, macrophages were incubated with equal numbers of DiD-labelled EVs at a final concentration of 4 × 10⁸ particles ml⁻¹ for 12 h. The culture medium was then removed, and cells were fixed with 4% formaldehyde for 15 min. After three PBS washes, cells were permeabilized with 0.1% Triton X-100 for 5 min and washed three additional times with PBS. Nuclei were counterstained with 0.5 μg ml⁻¹ 4′,6-diamidino-2-phenylindole (DAPI; Sigma-Aldrich) for 5 min. Fluorescence images were acquired using a confocal laser-scanning microscope.

#### 5.4.3 Proteomic profiling of EVs

##### 5.4.3.1 Sample Preparation and Protein Processing

Lysis was performed by adding 200 μL of lysis buffer (6 M urea [Sigma-Aldrich, Cat #U1250], 2 M thiourea [Sigma-Aldrich, Cat #T8656]) to each sample. After homogenization and centrifugation, proteins were quantified using a BCA assay (Thermo Scientific) after incubation at 37 °C for 30 min at 600 rpm.

For reduction and alkylation, add 30 μL lysis buffer to the protein solution, then add 5 μL 0.2 M Tris(2-carboxyethyl)phosphine (TCEP, Adamas-beta, Cat #61820E) and 2.5 μL 0.8 M iodoacetamide (IAA, Sigma-Aldrich, Cat #I6125). The samples were processed using the Barocycler system at 45 kpsi, with 30 s of high pressure and 10 s of ambient pressure, for 90 cycles at 30 °C.

##### 5.4.3.2 Enzymatic Digestion

Enzymatic digestion was carried out using Trypsin (Trypsin Gold, MS-grade, Hualishi Tech, Cat #HLS TRY001C) and recombinant Lys-C (rLys-C, MS-grade, Hualishi Tech, Cat #HLS LYS001C). Trypsin was reconstituted in 1 mM HCl or 50 mM acetic acid (final concentration 0.5 μg/μL). Each sample was supplemented with 75 μL of 0.1 M ABB, 5 μg rLys-C, and 1.25 μg trypsin, and then digested using the Barocycler at 20 kpsi, with 10 s of high pressure followed by 10 s of ambient pressure, for 120 cycles at 30 °C.

After digestion, add 15 μL of 10% trifluoroacetic acid (TFA, Thermo Fisher Scientific, Cat #85183) to terminate the reaction. Adjust the final pH to 2–3.

##### 5.4.3.3 Desalting

Peptide desalting was performed using SOLAμ SPE plates (Thermo Fisher Scientific™, San Jose, USA). The cartridges were activated with 200 μL of 100% methanol, equilibrated with 200 μL of 0.1% TFA in water, and washed twice with 2% ACN containing 0.1% TFA. Samples were loaded in 200 μL of 2% ACN with 0.1% TFA, washed again, and eluted with 100 μL of 40% ACN with 0.1% TFA. Peptides were dried by vacuum centrifugation and resuspended in 0.1% formic acid before LC-MS/MS analysis.

##### 5.4.3.4 Proteome Sample Analysis

Liquid chromatography-mass spectrometry (LC-MS) analysis was performed on a Vanquish Neo UHPLC system coupled to an Orbitrap Astral mass spectrometer (Thermo Scientific, San Jose, USA) for data-independent acquisition (DIA). The mobile phase A consisted of 98% water, 0.1% formic acid, and 2% Acetonitrile. Mobile phase B comprised 20% water, 80% acetonitrile, and 0.1% formic acid. All reagents were of MS grade.

For the DIA acquisition, the peptide concentration was set to 0.1 μg/μL, with an injection volume of 4 μL. The amount of sample loaded for each DIA acquisition was 400 ng. During sample acquisition, peptides were loaded onto a pre-column (5 µm, 5 mm×300 µm i.d.) at 800 bar, then eluted onto an analytical column (1.9 µm, 120 Å, 150 mm×75 µm i.d.) at a flow rate of 500 nL/min. An effective LC gradient (8% to 40% mobile phase B) was used for analysis, with a run time of 18 minutes.

The mass spectrometry scan parameters were configured as follows: the FAIMS voltage was set to -42 V. The primary scan covered a mass-to-charge (m/z) range of 380 to 980, with a resolution of 240,000. The normalized AGC target was set to 500%, and the maximum injection time (max IT) was 3 ms. For secondary scans, the m/z range was adjusted to 150-2000, maintaining the normalized AGC target at 500%. The collision energy was calibrated to 25%, with a maximum injection time of 3 ms. The precursor ion mass range for these scans was set to 380-980 m/z, with an isolation window of 2 m/z and no window overlap. A total of 299 windows were utilized for the analysis.

##### 5.4.3.5 Mass Spectrometry Data Analysis

The mass spectrometry data were processed using the DIA-NN software (version 1.9.1) for database searching. Carbamidomethylation of cysteine was set as a static modification, while oxidation of methionine was set as a variable modification. This comprehensive analysis provided both qualitative and quantitative data, applying a stringent false discovery rate (FDR) threshold of 0.01 or lower as the filtering criterion.

### 5.5 Molecular biology experimental characterization

#### 5.5.1 Live/dead staining assay

The viability of MSCs in microcapsules was assessed using a LIVE/DEAD kit (Invitrogen, Shanghai, China) according to the manufacturer’s protocol. Specifically, 2 mM calcein-AM and ethidium homodimer-1 were added to a culture plate for 10 min at 37 °C. Microcapsules were washed three times with the medium and observed using a Confocal Laser Scanning Microscope (OLYMPUS FV3000, Japan).

#### 5.5.2 Cell proliferation

The proliferation of MSCs on 2D polyacrylamide hydrogels was evaluated using a Cell Counting Kit-8 (CCK8, Solarbio, China). A total of 2 × 10⁵ MSCs were seeded in polyacrylamide hydrogels with varying stiffness in 24-well plates and cultured for 24 hours and 72 hours, respectively. At the designated time points, 100 μL of the supernatant was transferred to 96-well culture plates, and 10 μL of CCK-8 solution was added to each well. After incubation at 37°C for 4 hours, the absorbance at 450 nm was measured using a microplate reader (Bio-Rad iMark, United States).

#### 5.5.3 Cell cytomorphology

Low-concentration cells (1 × 10⁴ MSCs) and high-concentration cells (2 × 10⁵ MSCs) were encapsulated in 200 μL polyacrylamide hydrogels with different stiffnesses and cultured for 1 and 3 days, respectively. For visualization of cell nuclei and F-actin, the samples were stained with 0.5 μg/mL 4’,6-diamidino-2-phenylindole (DAPI, Sigma-Aldrich, USA) for 5 minutes and phalloidin (A12379, Thermo Fisher Scientific, USA, 1:400 dilution) for 1 hour at room temperature. The samples were washed twice with phosphate-buffered saline (PBS, Servicebio, China) and then observed using a confocal laser scanning microscope (FV3000, Olympus, Japan).

#### 5.5.4 Quantitative real-time PCR

Total RNA was isolated from cultured cells and reverse-transcribed into first-strand cDNA using the NovoScript 1st Strand cDNA Synthesis Kit (Novoprotein, Suzhou, China; Cat. No. E041). Quantitative real-time PCR (qPCR) was performed using 2 × HyperMB Universal SYBR Green qPCR Master Mix (Sangon Biotech, Shanghai, China; Cat. No. B690016-0001) on a Bio-Rad CFX96 Real-Time PCR Detection System (Bio-Rad Laboratories, Hercules, CA, USA). β-actin was selected as a housekeeping gene.

#### 5.5.5 Western Blot Analysis

MSCs were lysed using RIPA buffer (containing 25 mM Tris-HCl at pH 7.6, 150 mM NaCl, 1% Triton X-100, 1% sodium deoxycholate, 1 mg/mL PMSF, and 0.1% SDS) for 30 minutes, while EVs were lysed using BeyoExo™ EV Lysis Buffer (Beyotime, C3632) for 30 minutes. Both sample sets were centrifuged at 12,000 rpm for 20 minutes, and the supernatants were collected. The MSC samples were incubated at 37°C for 45 minutes to denature proteins, whereas EV protein samples were incubated at 95°C for 10 minutes. A BCA Protein Assay Kit was used to generate a standard curve for protein quantification, allowing for the calculation of the specific loading volume for each sample. After electrophoresis and membrane transfer, proteins were transferred onto PVDF membranes and blocked with 5% non-fat milk for 2 hours. Membranes were subsequently incubated overnight at 4°C with primary antibodies specific for ABCA1 (Abcam; Cat. No. ab66217), CD9 (UpingBio, Shanghai, China; YP-mAb-16288), CD81 (UpingBio, Shanghai, China; YP-mAb-16294), TSG101 (UpingBio, Shanghai, China; YP-mAb-16293), Flotillin-2 (UpingBio, Shanghai, China; YP-mAb-03874), CD63 (UpingBio, Shanghai, China; YP-mAb-16296), Calnexin (UpingBio, Shanghai, China; YP-mAb-16285), GLOD4 (UpingBio, Shanghai, China; YP-mAb-10035), and β-actin (Proteintech, Wuhan, China; Cat. No. 66009-1-Ig), followed by incubation with a goat anti-rabbit IgG H&L (HRP)-conjugated secondary antibody for 1 hour at room temperature.

#### 5.5.6 Immunofluorescence staining

Cells were fixed with a 4% paraformaldehyde solution and rinsed three times with 1× PBS. They were then permeabilized with 0.1% Triton X-100 for 15 min at room temperature. 1 % bovine serum albumin (BSA) was then used to block for 20 min, and the cells were incubated with rabbit polyclonal to iNOS (Abcam, ab15323), Goat polyclonal to liver Arginase (Abcam, ab60176), YAP1 Polyclonal antibody (Proteintech,13584-1-AP), or Rabbit polyclonal to Collagen I (Abcam, ab34710) in blocking solution overnight at 4 °C. The cells were washed 3 times with PBS and then incubated with 1:200 Alexa Fluor 594 Goat anti-Rabbit IgG (H + L) Cross-Adsorbed Secondary (Invitrogen, A11012) or 1:200 Alexa Fluor 488 Donkey anti-Goat IgG (H + L) Cross-Adsorbed Secondary Antibody (Invitrogen, A11055) in 1 % BSA for 60 min at room temperature in the dark. Finally, DAPI (Sigma, USA) was used to stain the nuclei, and the samples were observed under a confocal laser microscope.

#### 5.5.7 ELISA and secretome analysis

The macrophage supernatant was centrifuged at 3,000 rpm for 20 minutes, and the levels of IL-1β, IL-6, IL-10, and TNF-α were measured using mouse ELISA kits (Thermo Fisher, Shanghai) following the manufacturer’s instructions. Bronchoalveolar lavage fluid (BALF) was centrifuged at 3,000 rpm for 20 minutes, and the supernatant was diluted 2-5 fold to ensure the sample concentration fell within the detection range of the kits. Inflammatory cytokines in the BALF (IL-4, IL-5, IL-9, IL-10, IL-12p40, RANTES, MIP-1α, MIP-1β, GCSF, GM-CSF, CCL11, TNF-α, IL-13, IL-1β, IL-6, IL-12p70, IL-17A, MCP-1, CXCL1, IFN-γ, IL-1α, IL-2, and IL-3) were assayed using the same ELISA method (UpingBio, China)

#### 5.5.8 Plasmid construction and transfection

Human GLOD4 overexpression plasmids and siRNAs targeting GLOD4 (si-GLOD4) were synthesized and amplified by Hippo Biotechnology Co., Ltd. Human siRNA targeting ABCA1 (si-ABCA1) was purchased from Sangon Biotech (Shanghai, China).

For ABCA1 knockdown experiments, human umbilical cord-derived MSCs were transfected with si-ABCA1 according to the manufacturer’s instructions. At 48 h after transfection, ABCA1 expression was assessed at the mRNA and protein levels by RT– qPCR and Western blotting, respectively.

For GLOD4 manipulation, immortalized human bone marrow-derived MSCs were transfected with either the GLOD4 overexpression plasmid or si-GLOD4. GLOD4 expression was similarly evaluated by RT–qPCR and Western blotting 48 h after transfection to confirm successful overexpression or knockdown.

For experiments requiring subsequent hydrogel-based mechanical conditioning, cells were collected 24 h after transfection, counted, and seeded at a fixed density of 1 × 10⁶ cells per condition. After overnight attachment, cells underwent the indicated hydrogel-based mechanical or mechanochemical treatments, followed by downstream EV collection and functional analyses.

### 5.6 Animal experiment

This animal study was conducted in strict compliance with the regulations of the Biomedical and Animal Ethics Committee of Dalian University of Technology (Approval No.: DUTSBE230905-01). Eight-week-old male C57/BL6 mice were used in the experiments. To establish the pulmonary fibrosis (PF) model, mice under light anesthesia received a single intratracheal injection of 1 mg/kg bleomycin sulfate (BLM, Solarbio, China, Cat. No. IB0871) solution. On day 7 post-injury, 8×10^8^ particles of EVs were delivered into the lungs via intratracheal injection, with mice injected with normal saline serving as the control group. At 1, 3, 5, 7, and 14 days after treatment with DID-labeled 8×10^8^ particles of EVs, the mice were euthanized, and lung tissues were harvested. The distribution of EVs in the lungs was visualized using an in vivo imaging system (LB 983 NC100, Germany). Animals were euthanized on day 21 post-BLM injury. After PBS perfusion to collect bronchoalveolar lavage fluid (BALF), lung tissues were obtained for subsequent analysis.

## Acknowledgement

The National Natural Science Foundation of China generously funded this work (No. 3271748, No. 82501251, No. 52403152, No. 52302344).

## AI declaration

We acknowledge the use of ChatGPT (https://chatgpt.com/) to assist with grammar checking and writing refinement. Prompts used included “check the following text for grammatical errors” and “revise the following text to improve concision and clarity without changing the scientific meaning”. All scientific content was reviewed and verified by the authors, who take full responsibility for the content.

## Data Availability

The data supporting the findings of this study are available from the corresponding author upon reasonable request.

## Author contributions

P.C. planned and executed the experiments, analysed the data, and was involved in discussions of the data. P.C., A.C. and W.H. wrote the paper. H.Z., C.K., H.Y., T.T., and W.X. performed the experiments. All authors critically reviewed and approved the paper.

## Declaration of competing interests

The authors declare that they have no known competing financial interests or personal 1012 relationships that could have appeared to influence the work reported in this paper.

## Notes

### Competing Interest Statement

The authors have declared no competing interest.

### Summary of Updates

The manuscript has been carefully revised in response to the reviewers' comments, and the updated preprint is now available on bioRxiv.

